# Long-read direct RNA sequencing of the mitochondrial transcriptome of *Saccharomyces cerevisiae* reveals condition-dependent intron turnover

**DOI:** 10.1101/2023.01.19.524680

**Authors:** Charlotte C. Koster, Askar Kleefeldt, Marcel van den Broek, Marijke Luttik, Jean-Marc Daran, Pascale Daran-Lapujade

## Abstract

Mitochondria fulfil many essential roles and have their own genome, which is expressed as polycistronic transcripts that undergo co- or post-transcriptional processing and splicing. Due to inherent complexity and limited technical accessibility of the mitochondrial transcriptome, fundamental questions regarding mitochondrial gene expression and splicing remain unresolved, even in the model eukaryote *Saccharomyces cerevisiae*. Long-read sequencing could address these fundamental questions. Therefore, a method for enrichment of mitochondrial RNA and sequencing using Nanopore technology was developed, enabling the resolution of splicing of polycistronic genes and the quantification the spliced RNA.

This method successfully captured the full mitochondrial transcriptome and resolved RNA splicing patterns with single-base resolution, and was applied to explore the transcriptome of *S. cerevisiae* grown with glucose or ethanol as sole carbon source, revealing the impact of growth conditions on mitochondrial RNA-expression and splicing. This study uncovered a remarkable difference in turn-over of group II introns between yeast grown in mostly fermentative and fully respiratory conditions. Whether this accumulation of introns in glucose medium has an impact on mitochondrial functions remains to be explored. Combined with the high tractability of the model yeast *S. cerevisiae*, the developed method enables to explore mitochondrial transcriptome regulation and processing in a broad range of conditions relevant in human context, including aging, apoptosis and mitochondrial diseases.

## Introduction

Mitochondria are one of the hallmarks of eukaryotic cells. Commonly known as ‘the powerhouse of the cell’, they perform several key cellular processes such as iron-sulfur clusters, branched amino acids, heme, and lipids biosynthesis and play an important role in cellular aging and programmed cell death (reviewed by (1,2)). To date, about 150 mitochondrial disorders have been reported in humans, which genetic origin lies in mitochondrial (mtDNA) or nuclear (nDNA) DNA. As remnant of their bacterial origin, mitochondria harbor their own DNA, however most of the 1.000 – 1.500 proteins composing mitochondria are nuclearly-encoded and imported from the cytosol to mitochondria (3). Remarkably, mitochondria have retained specific genes and require their own complete set of proteins for gene expression, RNA splicing and degradation and DNA maintenance (4). Mitochondrial functions are largely conserved across eukaryotes, including the eukaryotic model *Saccharomyces cerevisiae*, which is a preferred model for mitochondrial biology for two main reasons. Firstly, *S. cerevisiae* can grow in the complete absence of respiration and with partial or complete loss of mtDNA (5). Secondly, *S. cerevisiae* is still to date one of the rare organisms harboring genetic tools to modify mtDNA beyond base editing (*e*.*g*. gene deletion, integration (6)). Using these features, mtDNA-free yeast cells (ρ^0^ mutants) were shown to stably maintain mouse mtDNA (7), a first step towards the potential humanization and engineering of mtDNA in yeast. *S. cerevisiae* has therefore an important role to play in deepening our understanding of mitochondrial processes.

The *circa* 86 kb *S. cerevisiae* mitochondrial genome encodes seven oxidative phosphorylation proteins (*COX1, COX2, COX3, COB, ATP6, ATP8, Q0255*), two ribosomal subunits (15s rRNA and 21s rRNA), a ribosomal protein (*VAR1*), the RNA subunit of RnaseP (*RPM1*), and a full set of 24 tRNAs (8). Unlike its nDNA, *S. cerevisiae* mtDNA has a prokaryote-like structure with 11 polycistronic primary transcripts (Figure 1), which expression is regulated by T7-like promoters (9,10). Conserved dodecamer sequences separate the transcripts within the polycistrons and enable modulation of gene expression (11-13). While splicing is rare in *S. cerevisiae* nDNA (14), most mitochondrial genes (*COX1, COB, Q0255, VAR1* and *21srRNA*) contain intronic sequences (8). Several of these introns encode proteins, *e*.*g*. the I-SceI homing endonuclease is encoded in the intron of the 21s rRNA gene (15). Furthermore, because of alternative splicing, parts of an exon combined with different intronic sequences of the same primary transcript yield different proteins, next to the exon-encoded protein. These splicing events are catalyzed by nDNA- and mtDNA-encoded enzymes (16). Despite its small size, the mtDNA results in a complex RNA landscape involving mechanisms that, despite decades of study, are not fully elucidated.

**Figure 1.**
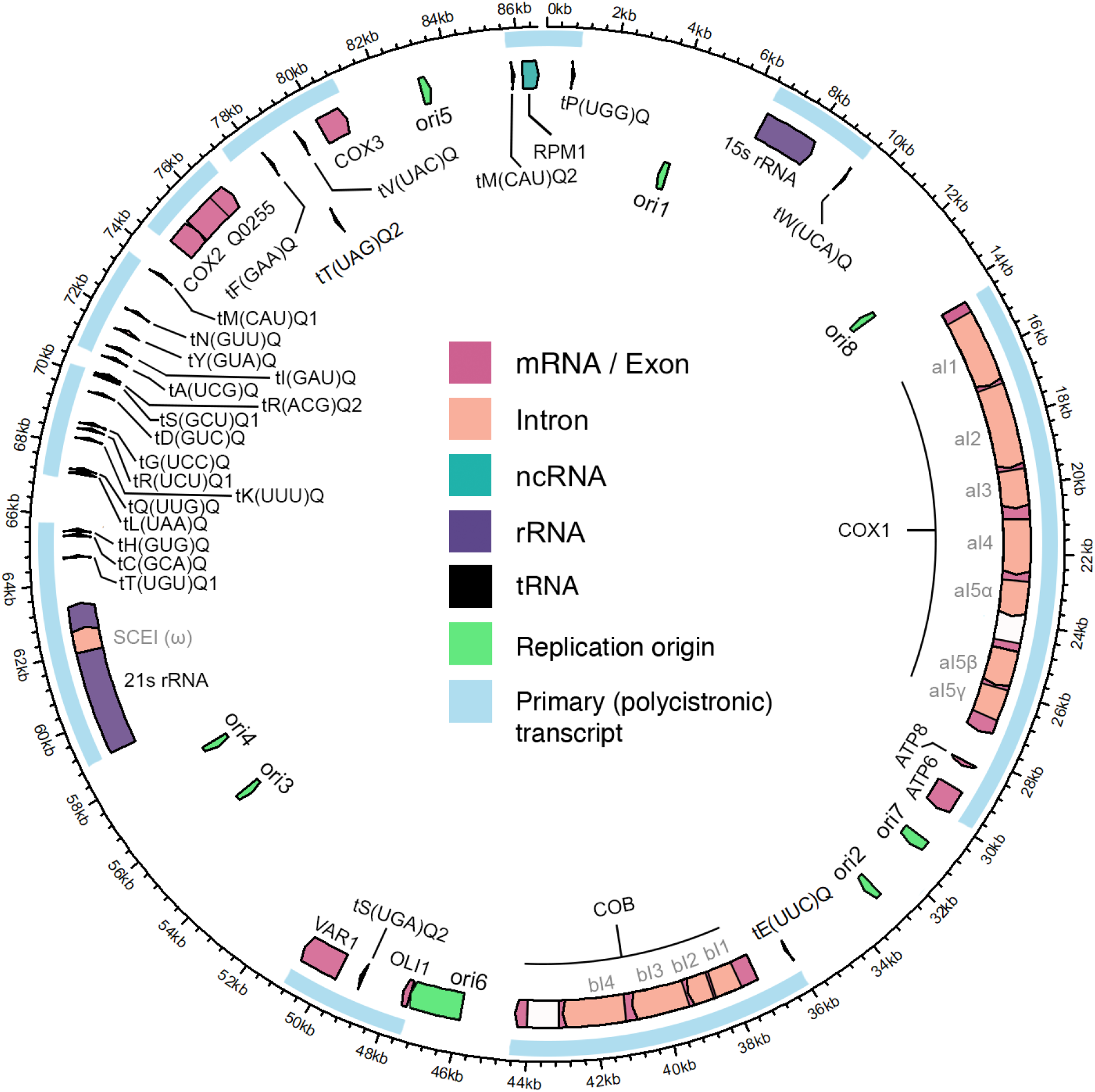
Annotated mitochondrial genome of CEN.PK113-7D. Primary polycistronic transcripts (light blue boxes); mRNA: messenger RNA, ncRNA: noncoding RNA, rRNA: ribosomal RNA, tRNA: transfer RNA. Exons that encode the main proteins are indicated in pink. Coding intron sequences are encoded in orange, non-doing intron sequences are indicated in white.

Since the first gene arrays in the ‘90s, the cytosolic transcriptome of *S. cerevisiae* has been intensively investigated (17,18). The mitochondrial transcriptome is generally not detectable in these classical transcriptome studies, which is largely explained by the fact that mRNA extraction methods rely on selection of poly(A)-tailed mRNA, while yeast mtRNA is not polyadenylated (19). Additionally, the mitochondrial transcriptome represents a small fraction of the total yeast transcriptome (ca. 5 %), which is largely dominated by (ribosomal) cytosolic RNA species (20,21). Capturing the mitochondrial transcriptome therefore requires a dedicated methodology. To date, a single study reported the yeast mitochondrial transcriptome, using cellular subfractionation to enrich for mitochondria and short read RNA sequencing (13). While this study brought new insight in regulatory elements such as promoter sequences, dodecamer sequences, UTR’s and processing sites, the use of short reads, even with paired ends, cannot capture the full complexity of the mtRNA transcriptome. This technical limitation can be overcome by implementing long-read RNA sequencing, as recently demonstrated by the successful resolution of various complex and spliced transcriptomes ranging from viruses to plant, mammalian and even a full poly(A)-human transcriptome (18,22-27).

The goal of this study is to provide a comprehensive description of the mitochondrial transcriptome of *Saccharomyces cerevisiae*, including identification of processing of polycistronic transcripts and splicing of the different transcript isoforms. To this end, we developed a robust protocol for mitochondrial RNA isolation, and combined it with Nanopore long-read direct RNA sequencing technology. This method was used to investigate the response of the mitochondrial transcriptome to different carbon sources (glucose and ethanol) chosen for their ability to tune yeast physiology and respiratory activity.

## Materials and Methods

### 1. Strains & culture conditions

*Saccharomyces cerevisiae* strains used in this study (Table 1, Table S 1) were derived from the CEN.PK lineage (28), or from a 161 lineage (also known as ID41-6/161) (29-31), which were a gift from prof. Alan Lambowitz (University of Texas at Austin, Austin, TX, USA).

**Table 1.** *S. cerevisiae* strains used in this study.

| Strain name | Relevant genotype | Parental strain | Reference |
| --- | --- | --- | --- |
| CEN.PK113-7D | *MATα* | - | Entian and Kötter (28) |
| CEN.PK113-5D | *MATα ura3-52* | - | Entian and Kötter (28) |
| 161-U7 / 1<sup>+</sup>2<sup>+</sup> | *MATa ade1 lys1 ura3*, no mitochondrial ω-intron | - | Mahler and Lin (29) |
| 161-U7 I<sup>0</sup> | *MATa ade1 lys1 ura3*, no mitochondrial ω-intron, no introns in *COX1* and *COB* | 161-U7 / 1<sup>+</sup>2<sup>+</sup> | Huang *et al.* (31) |
| IMC251 | *MATα ura3-52* + pUDC329 (*preCOX4-mTq2, URA3*) | CEN.PK113-5D | This study |
| IMC252 | *MATa ade1 lys1 ura3*, no mitochondrial ω-intron + pUDC329 (*preCOX4-ymTq2, URA3*) | 161-U7 / 1<sup>+</sup>2<sup>+</sup> | This study |
| IMC253 | *MATa ade1 lys1 ura3*, no mitochondrial ω-intron, no introns in *COX1* and *COB* + pUDC329 (*preCOX4-ymTq2, URA3*) | 161-U7 I<sup>0</sup> | This study |

Yeast strains were grown aerobically at 30 °C in 500 mL shake flasks containing 100 mL synthetic medium with ammonium as nitrogen source (SM) supplied with vitamins and trace elements, prepared and sterilized as described previously (32), in Innova incubator shakers (Eppendorf, Hamburg, Germany) set at 200 rpm. For respiro-fermentative growth sterilized media were supplemented with a D-glucose solution to a final concentration of 20 g L^-1^ glucose (SMD, ‘glucose media’). For respiratory growth, media were supplemented with absolute ethanol to a final concentration of 2 % (v/v) and with a glycerol solution to a final concentration of 2 % (v/v) (SMEG, ‘ethanol media’). Glucose solutions (50 % w/v) and glycerol solutions (99 % w/v) were autoclaved separately for 10 minutes at 110 °C. When required, medium was supplemented with separately sterilized solutions of uracil (Ura) to a final concentration of 150 mg L^-1^, Lysine (Lys) to a final concentration of 300 mg L^-1^ and/or Adenine (Ade) to a final concentration of 200 mg L^-1^ (33). Alternatively, YPD medium, containing 10g L^−1^ Bacto Yeast extract, 20 g L^−1^ Bacto Peptone and 20 g L^-1^ glucose was used.

For plate cultivation 2% (w/v) agar was added to the medium prior to heat sterilization.

Frozen stocks were prepared by addition of sterile glycerol (30% v/v) to late exponential phase shake-flask cultures of *S. cerevisiae* and 1 mL aliquots were stored aseptically at −80 °C.

### 2. Molecular biology & strain construction

Strains expressing the yeast-optimized fluorescent protein ymTurquoise2 (*ymTq2*, (34)) in the mitochondria (IMC251, IMC252 and IMC253, Table 1) were transformed with plasmid pUDC329 (*pTEF1-preSU9-ymTq2-tENO2, URA3, CEN/ARS, bla, ColE1)*. Plasmid pUDC329 was cloned using Golden-Gate assembly performed according to Lee *et al*. (35), from part plasmids pUD538 (GFP-dropout backbone, *URA3, CEN/ARS, AmpR, ori)*, pUD585 (part 2, *pTEF1*), pGGkp320 (part 3a, *preSU9*), pGGkp308 (part 3b, *ymTurquoise2*), pYTK055 (part 4, *tENO2*) (Table S 2)(36,37). The part plasmids were either obtained from the Yeast Toolkit collection (35) or made according to the author’s instructions using primers with Golden Gate flanking regions corresponding to the respective part type. The *preSU9* sequence was ordered as a synthetic sequence at GeneArt (Supplementary Information, Life technologies, Carlsbad, CA). pDRF1-GW ymTurquoise2, *pFA6a-link-ymNeongreen-URA3* and pDRF1-GW ymYPET were a gift from Bas Teusink (Addgene plasmid # 118453, #125703, # 118455; (34)). 1 μL of the Golden-Gate reaction mixture was transformed into *E. coli* cells (XL1-Blue, Agilent Technologies, Santa Clara, CA), which were grown in solid Lysogeny broth (LB) medium (5.0 g L^−1^ yeast extract, 10 g L^−1^ Bacto tryptone, 2 % Bacto agar (BD Biosciences, Franklin Lakes, NJ), 5.0 g L^−1^ NaCl) supplemented with 100 mg L^−1^ ampicillin. Upon growth, random colonies were picked and resuspended in 10 μL sterile water, of which 1 μL was used for diagnostic colony PCR, using using DreamTaq MasterMix 2X (Thermo Fisher Scientific, Waltham, MA)) and desalted primers (Sigma-Aldrich, St. Louis, MO, Table S 3), with an initial 10-minute incubation at 95 °C to release *E. coli DNA*. Positive transformants were inoculated in 5 mL liquid LB medium (omitting the Bacto agar) and plasmids were isolated using a GeneJET Plasmid Miniprep Kit (Thermo Fisher Scientific). Correct plasmid structure was confirmed using restriction analysis using the FastDigest KpnI enzyme (Thermo Fisher Scientific) according to the manufacturer’s instructions and by EZseq Sanger sequencing (Macrogen, Amsterdam, the Netherlands). Fragment sizes were assessed on a 1 % agarose gel. 100 ng plasmid was transformed into *S. cerevisiae* cells as previously described (38), mutants were selected on SMD (+Lys/Ade) medium lacking uracil. Successful transformation with the plasmid was confirmed using fluorescence microscopy, mutants were re-streaked three times on selective solid medium to obtain single colony isolates, which were grown and stored as described before. Construction of strain IMC173, used for optimization of the mitochondrial isolation protocol (Table S 1,Figure S 1) is described in the Supplementary Information (39,40).

### 3. Preparation of mitochondrial RNA & sequencing

#### Isolation of mitochondria by differential centrifugation

Crude mitochondria were isolated by a combination of protocols described by (41) and (42), with a few modifications. The protocol was initially tested using a strain (IMC173) with fluorescent mitochondria prior to isolating mitochondria for RNA-seq (Figure S 1, Supplementary methods). Pre-cultures of CEN.PK113-7D were grown overnight on SMD and were used to inoculate duplicate 500 mL shake flasks containing 100 mL SMD or SMEG at an optical density at 660 nm (OD_660_) of 0.1 - 0.2. The mitochondrial volume of a cell is dependent on respiratory activity (43), therefore, to yield enough mitochondrial mass, 300 mL of culture were used when grown on respiratory SMEG media and 800 mL of culture for respiro-fermentative glucose media. SMD cultures were grown until mid-exponential phase (OD_660_ of 5) to prevent cells from going through diauxic shift. SMEG cultures were grown until mid- to late-exponential phase (OD_660_ of 5-19), before harvesting by centrifugation. All centrifugation steps were performed at 4 °C in an Avanti J-E high-speed fixed-angle centrifuge (Beckman Coulter Life Sciences, Indianapolis, IN) with a JA-10 rotor for volumes larger than 30 mL and a JA-25.50 rotor for 30 mL and less. Stock solutions were used to prepare working solutions and were sterilized prior to use at 121 °C for 30 minutes. The following stock solutions were prepared: potassium phosphate buffer pH 7.4 (30.4 mM KH_2_PO_4_, 69.6 mM K_2_HPO_4_), Tris buffer (0.1 M Tris, pH 7.4), HEPES-KOH (0.5 M HEPES, pH set to 7.2 with KOH), MgCl·6H_2_O (0.1 M). A sorbitol stock solution (3 M) was autoclaved at 110 °C for 10 minutes. Working solutions were freshly prepared prior to use.

Cultures were centrifuged in pre-weighed bottles for 5 min at 1,500 × *g* to remove media, then washed with Tris (0.1 M Tris, pH 7.5) and centrifuged again. The supernatant was thoroughly removed, the weight of each pellet was determined, and pellets were resuspended in 30 mL Tris-DTT (0.1 M Tris, 10 mM DTT, prepared fresh before use, pH 9.3) and transferred to centrifuge tubes. Cultures were incubated in Tris-DTT for 15 minutes in a waterbath at 30 °C with gentle shaking, then centrifuged 10 min at 4,342 × *g*, washed once in Sorbitol-Phosphate buffer (SP-buffer, 1.2 M sorbitol, 20 mM potassium phosphate buffer pH 7.4) and resuspended in SP-buffer. Zymolyase 20T (AMSbio, Alkmaar, the Netherlands) was added to each culture at a concentration of 5 – 10 mg per gram pellet wet weight. Cultures were incubated in presence of Zymolyase in a waterbath at 30 °C with gentle shaking to obtain spheroplasts. Spheroplasting progress was monitored by resuspending 10 μL of spheroplasts in 2 mL of dH_2_O and measuring the OD_660_, as the low osmotic pressure of water will cause bursting of spheroplasts. Spheroplasting was halted after the OD_660_ dropped by 50 - 75 %, (ca. 30 – 45 min). Spheroplasts were always kept on ice.

Spheroplasts were pelleted 10 min at 4,342 × *g*, and supernatant containing Zymolyase was thoroughly removed. Spheroplasts were then resuspended in 20 mL SP-buffer by gentle shaking and pelleted again (10 min at 4,342 × *g*). The washed and pelleted spheroplasts were resuspended in low-osmolarity Sorbitol-HEPES buffer (SHE, 0.6 M sorbitol, 20 mM HEPES-KOH, 2 mM MgCl) with with cOmplete Protease Inhibitors Cocktail (PI, Roche Diagnostics, Rotkreuz, Switzerland) and homogenized at 4 °C using a pre-cooled Potter-Elvehjem PTFE pestle at 100 RPM and glass tube with a working volume of 30 mL. The homogenate was cleared of cellular debris by centrifugation at 1,500 × *g* for 10 min. The supernatant was collected with a pipette, transferred to a clean tube, and centrifuged at 12,000 × *g*. The supernatant containing the cytosolic fraction was collected in a separate tube. The pellet, containing the mitochondria-enriched fraction was resuspended in 2 mL of SHE + PI and immediately processed for RNA extraction or flash-frozen in liquid nitrogen and stored at -80 °C.

#### RNA extraction from mitochondrial fraction

RNA-extraction and handling was done in an RNase-free workspace. When possible, materials and workspaces were cleaned with RNAseZAP spray and towels (Sigma Aldrich, St. Louis, MO) and consumables and working solutions were autoclaved for 45 minutes at 121 °C to denature any RNases. *In vitro* synthesized RNA was spiked throughout the protocol to assess throughput and efficiency of the protocol. The synthesis of control RNA and timing of spike-ins is described in the supplementary methods.

RNA from the mitochondria-enriched fraction was isolated using the miRNeasy Mini kit (Qiagen, Venlo, the Netherlands). Mitochondrial fractions were spun down for 10 minutes 10,000 × *g* at 4 °C, the buffer was removed, and the mitochondria were thoroughly resuspended in 700 μL QIAzol reagent at RT by vortexing. This was sufficient to lyse the mitochondria, and RNA was extracted following manufacturer’s instruction. On-column DNase I treatment using the RNase-free DNase set (Qiagen) was included in the RNA extraction to remove any DNA contaminants. RNA was eluted in 30 μL nuclease-free water at RT. RNA integrity (RIN) was assessed using an RNA ScreenTape assay for the Tapestation (Agilent, Santa Clara, CA), and RNA quantity using a Qubit fluorometer with the RNA broad range assay kit (Thermo Fisher, Waltham, MA). Samples with an RIN > 7 and a yield of 3 μg or more were used for enzymatic polyadenylation.

#### *In vitro* polyadenylation and cleanup of isolated RNA

*In vitro* polyadenylation was performed using *E. coli* Poly(A) Polymerase (New England Biolabs, Ipswich, MA), according to manufacturer’s instructions, with a few modifications: 3 – 10 μg of RNA was added to each reaction to meet Nanopore input requirements. Additionally, 20 U murine RNase inhibitor (New England Biolabs) were added to each reaction. After polyadenylation, samples were cleaned up miRNeasy Mini kit with a few modifications. The polyadenylation reaction was filled up to 100 μL using nuclease-free water. Then 350 μL RLT buffer (RNeasy kit, Qiagen) were added to the sample to replace QIAzol buffer. Next, 250 μL 100 % Ethanol were added and the whole volume was transferred to a miRNeasy spin column. Column purification was further performed according to manufacturer’s instructions, omitting the on-column DNase digestion. The RNA was eluted in 30 μL nuclease-free water and RIN was determined with an RNA Screentape assay, and quantity with a Qubit fluorometric assay. Presence of contaminants was determined by measuring absorbance using a Nanodrop spectrophotometer (Thermo Fisher). Purified RNA with an RIN of > 7, a concentration of > 55 ng/μL, a 260/280 absorbance ratio of around 2.0 and a 260/230 ratio between 2.0 and 2.2 was deemed high quality and used for RNA sequencing.

#### *In vitro* synthesis of RNA spike-in controls

RNA spike-in control sequences were generated by in vitro transcription (IVT) of DNA templates bearing a long T7 promoter. Three sequences were of 60 bp length (IVT01, IVT02, IVT03), one of 102 bp sequence (IVT04), one 729 bp sequence (IVT-Neon), one 742 bp sequence (IVT-YPet, (44)) and the 1026 bp IVT-eYFP sequence (IVT-YFP) (See: “synthetic RNA sequences used in this study”). DNA templates for 60 bp sequences (IVT01, IVT02 and IVT03) were generated by annealing of primers (Table S 2) by heating equimolar amounts of both primers up to 95°C for 5 minutes and cooling down slowly at room temperature for primers to hybridize. DNA template for IVT04 was generated by overlap PCR, where self-dimerizing primers were used in final concentrations of 0.2 μM in PCR reactions of 50 μl reaction volume using Phusion High fidelity polymerase (Thermo Fisher) according to manufacturer’s instructions. DNA templates for IVT-eYFP, IVT-Neon and IVT-YPet were generated by PCR amplification from plasmids using Phusion HF polymerase (Table S 2, Table S 4). All DNA templates were subject to PCR purification and size validated by agarose gel-electrophoreses. 0.5 – 1 μg DNA template were in vitro transcribed using HiScribe® T7 Quick High Yield RNA Synthesis Kit (New England Biolabs, Ipswich, MA), following manufacturer’s instruction: IVT reactions with DNA templates < 300 bp were performed in reaction volume of 20 μl and reactions were incubated at 37°C in a thermocycler for 4 – 16 hours. For DNA templates > 300 bp, the reaction volume was 30 μl, and incubation period was adjusted to 2 hours for 37°C in a thermocycler. IVT products were recovered directly after reaction by miRNeasy Mini Kit for products < 300 bp, where reactions were filled to 100 μl with nuclease-free water and buffer RLT from RNeasy Mini Kit (Qiagen, Venlo, the Netherlands) was added, omitting QIAzol. Alternatively, IVT products < 300 bp were purified by lithium chloride (LiCl) precipitation as described in manufacturer’s instructions. For IVT products >300 bp, RNeasy Mini Kit (Qiagen) was used, following manufacturer’s instructions. All IVT products were eluted in 60 μl nuclease-free water. Quantity and purity were assessed photospectrometrically by Nanodrop and Qubit. IVT product sizes were validated by denaturing gel-electrophoresis with denaturing 2% (w/v) agarose with 3% (v/v) formaldehyde gels in MOPS buffer (40 mM MOPS pH7, 10 mM Sodium Acetate, 1mM EDTA). Prior to loading on gel, 2X loading dye (NEB) was added the RNA and the sample was denatured at 70°C for 10 minutes. Samples were directly put on ice for 2 minutes, and subsequently loaded onto the gel. Synthetic RNA controls were spiked in at a final concentration of 45 fmol at two different timepoints: i) after isolation of the mitochondrial fraction, prior to RNA isolation, to simulate the presence of cytosolic RNA and ii) prior to polyadenylation, to assess the throughput and efficiency of polyadenylation, cleanup, and library preparation. Lastly, the ONT Direct RNA sequencing kit includes a synthetic *ENO2* control RNA, which was spiked during library preparation according to manufacturer’s instructions.

#### Library preparation & sequencing

Poly(A)-tailed RNA samples were used for library preparation using the Direct RNA Sequencing Kit (SQK-RNA002, Oxford Nanopore Technologies (ONT), Oxford, United Kingdom), following manufacturer’s instructions. Library preparation yield was measured with Qubit® 2.0 fluorometer and corresponding Qubit™ dsDNA BR Assay Kit. Prior to library loading, remaining active pores were measured by a flow cell check with MinKNOW software (ONT). Library was loaded onto a R.9.4.1 flow cell with FLO-MIN106D chemistry using the Flow Cell Priming Kit (EXP-FLP002, ONT) according to manufacturer’s instructions. Sequencing runs were started via MinKNOW with default parameters for 16 hours.

### 4. Computational methods for RNA sequencing data analysis

#### Availability of data

When indicated, scripts and files required for analysis and interpretation of the data mentioned hereafter are deposited on our research group’s servers and can be freely accessed at https://gitlab.tudelft.nl/charlottekoste/mitornaseq. For R-based analysis, RStudio (RStudio, Boston, MA, USA) was used. Raw RNA-seq data and normalized expression levels were deposited at NCBI (www.ncbi.nlm.nih.gov) GEO under accession number GSE219013.

#### Basecalling

Raw FAST5 files were basecalled on a high-performance computing cluster (GPU) using Guppy 4.4.1 (ONT) with default parameters for flow cell type and sequencing kit. Basecalled reads were categorized as PASSED for Guppy-determined quality score > 7 and as FAILED for reads < 7 (Table S 5). All PASSED and FAILED FASTQ files were concatenated into a single FASTQ file each and filtered by length to cut-off at 40 bp Basecalled reads were processed according to standard EPI2ME workflow (Metrichor Ltd., ONT, Oxford, UK) for FASTQ RNA control reads to assess sequencing accuracy and coverage of The RNA control strand (RCS).

#### Annotation of the CEN.PK113-7D mitochondrial genome

As reference for alignment, *S. cerevisiae* CEN.PK113-7D complete genome sequence ((45,46)) was used. The mitochondrial genome of CEN.PK113-7D was annotated by aligning nucleotide sequences of the S288c mitochondrial mapping data from the mitochondrial genome deposited in the *Saccharomyces* Genome Database (SGD: https://www.yeastgenome.org/ (47,48)) as well as the s288c mitochondrial genome annotated by Turk *et al*. (13) to the CEN.PK113-7D mitochondrial genome sequence using SnapGene software (http://www.snapgene.com, GSL Biotech, San Diego, CA). The resulting annotated mitochondrial CEN.PK113-7D genome (Figure 1) was exported as GenBank and annotation (.gff) file and is deposited at Gitlab. The map of the CEN.PK113-7D mitochondrial genome was visualized in R using the circlize package (file: *mtDNA_plot*.*Rmd*) (49).

#### Mapping

To account for control sequences, the FASTA file of the genome reference sequence was amended with FASTA files of all *in vitro* synthesized control sequences including the sequence of RCS (*ENO2* gene as provided by ONT). The resulting FASTA file and corresponding annotation file were used for mapping with minimap2 (50) (parameters: *-ax splice -uf -k14 --secondary=no*) to obtain a Sequence Alignment/Map (SAM) file (51). All *PASSED* reads longer 40 bp were aligned to the reference FASTA file. SAM files were filtered and converted to Binary Alignment/Map (BAM) files and indexed using SAMtools, whereby only uniquely aligning reads were extracted based on FLAGs 0 and 16.

Poly(A)-tail length was estimated by nanopolish-polya (25), for both *PASSED* and *FAILED* reads of one of the sequencing runs (Ethanol replicate #3), following provided instructions. For assessment and visualization of poly(A)-tail lengths of datasets, only poly(A)-tail estimations with nanopolish-polya specific QC-tag PASS were used.

#### Feature quantification and normalization

Annotation of alignment and quantification of features was performed using FeatureCounts (52), where above named reference file was supplied together with filtered BAM file with uniquely aligned reads. Importantly, long-read specific, overlap allowed, directional counting in terms of strandedness and fractional counting (parameters: *-L -O -s 1 --fraction*) were employed. The present datasets originated from three biological culture replicates, mitochondria isolation rounds and sequencing runs, leading to different read distributions and library sizes (Figure 2B, Table S 5). Quantitative analysis of expression levels therefore required library normalization. RPKM (Reads Per Kilobase Million) is the standard normalization method for RNA-seq, developed for short-read RNA sequencing in which read length (*ca*. 150 - 200 bp) is substantially shorter than gene length (e.g. several kb in yeast). RPKM normalizes to library size, but also to gene length, as to compensate for this direct correlation between read number and gene length. However, normalization by gene length is not applicable to long-read RNA sequencing as read length often approaches or even exceeds the gene length. The datasets were therefore only normalized over library depth in Counts-per-Million (CPM) in R (script: *readcounts*.*rmd*) using edgeR (53). For all quantification analysis, reads mapping to both the *S. cerevisiae* native *ENO2* feature as well as RCS feature, were considered to be control reads, as the two could not be distinguished based on sequence identity. Normalization between Ethanol and Glucose datasets was done by normalization over the mitochondrial dry weight (mtDW, R-script: *normalization_etoh-gluc*.*rmd*). To normalize, the input amount of cells in OD units was first normalized to cell dry weight (CDW) using the following relations (experimentally determined as described by Verduyn *et al*. (32) using 30 samples of different OD_660_):

**Figure 2.**
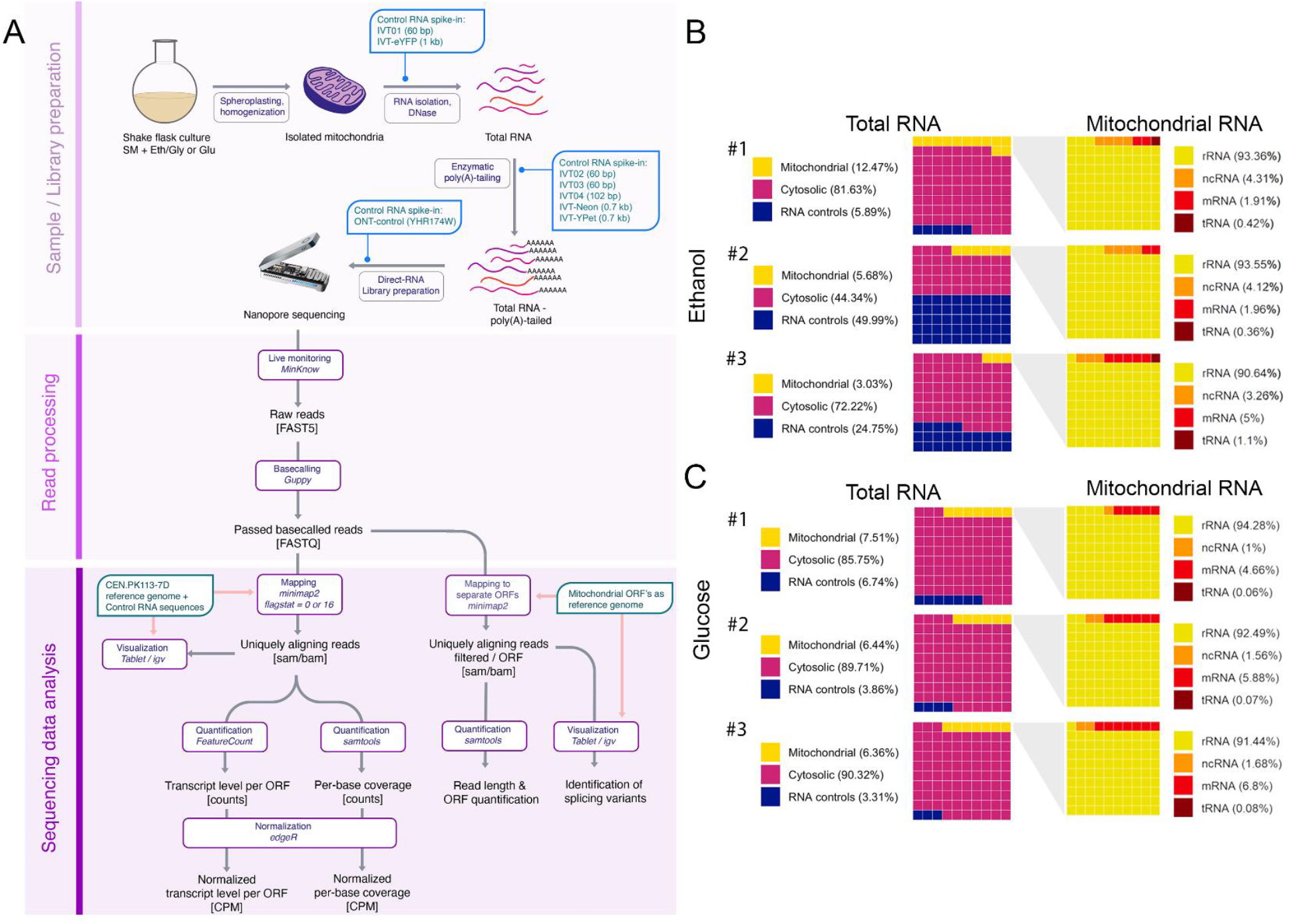
Sequencing pipeline and quality control. A) schematic overview of the sample preparation and sequencing pipeline. B,C) Waffle plots showing distribution of read origin and type of the passed reads (Q >=7) of the three replicates of ethanol-grown cultures (B) and glucose-grown cultures (C). The distribution of the origin of sequencing reads between mitochondrial, cytosolic and control RNA is shown on the left, on the right the distribution of different types of mitochondrial RNA is shown. One square represents 1/100 of the total number of reads.

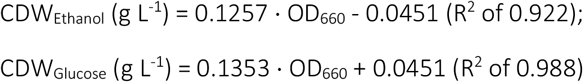

The CDW was multiplied by the volumes used (Table S 5), to get the total CDW per condition. Cells have a density of 1 g mL^-1^, and on ethanol a mitochondrial volume of 0.35 mL mitochondria per mL ethanol-grown culture and 0.05 mL mitochondria per mL glucose-grown culture (43). From this follows approximately 0.26 g (± 0.02) mtDW · g CDW^-1^ for ethanol-cultures and 0.027 (± 0.001) g mtDW · g CDW^-1^ for glucose cultures (± 0.001). This conversion factor was used to normalize over mitochondrial dry weight. All counts, either raw, normalized to CPM or normalized to CPM and mtDW are listed in the Supplementary Information (SI_readcounts.xlsx).

Per-base-coverage of the mitochondrial transcriptome was obtained with SAMtools (parameters: *samtools depth -aa*). Relative per-base-coverage at distinct loci were computed in R (script: *coverageplots*.*rmd*) by normalizing coverage at each base position to the total read depth in counts-per-million (CPM). Ratio’s between intron and exon levels were determined by calculating the coverage of the last 250 base pairs of each splicing variant or exon, as these base pairs are unique between each variant, and then subsequently normalizing over the exon level.

#### Read length estimation

For assessment of the read length estimation, a different reference alignment file was used in which the mitochondrial genome was split up into shorter reference sequences that each contained a single mitochondrial open reading frame (ORFs). This allowed for assigning of reads to single ORFs rather than to the full mtDNA, making quantification more straightforward and filtering out erroneous reads with gaps that span multiple ORFs (as shown in Figure S 11). Read mapping was then performed using the same parameters as described. Using the resulting BAM files, for each ORF the average length of its respective mapped reads was calculated (script: *readlengths*.*rmd*).

#### Visualization of read mapping

Read mapping was visualized using Integrated Genome Viewer (54) and Tablet (55). Mapping of long reads was visualized using the full mitochondrial genome as a reference sequence and corresponding BAM files. Splicing was visualized using separate mitochondrial ORFs as reference sequences with the respective BAM files, as this increases resolution of the data when looking into splicing events.

### 5. Characterization of intron-less strains

#### Fluorescence microscopy

Phase-contrast microscopy was performed using a Zeiss Axio Imager Z1 (Carl Zeiss AG, Oberkochen, Germany) equipped with a HAL 100 Halogen illuminator, HBO 100 illuminating system and AxioCam HRm Rev3 detector (60 N-C 1’’ 1.0×) (Carl Zeiss AG). The lateral magnification objective 100×/1.3 oil was used with Immersol 518F type F immersion oil (Carls Zeiss AG). Fluoresence of ymTurquoise2 was detected using filter set 47 (Carl Zeiss AG; excitation bandpass filter 436/20 nm, beamsplitter filter 455 nm, emission filter 480/40 nm). Results were analysed using the Fiji package of ImageJ (56). Mean fluorescence per cell was analyzed by thresholding the brightfield channel of each image using the Otsu algorithm to determine the location of the cells in the image. Based on the thresholded brightfield image, a binary mask was generated indicating the regions of interest (ROI, i.e. cell locations) of the brightfield image. Subsequently, the mask was overlaid on the DAPI channel of the image and the mean fluorescence for each ROI was determined, resulting in a mean fluorescence per cell.

#### Growth rate analysis

Growth rate analysis was performed in 96-wells microtiter plates at 30 °C and 250 rpm using a Growth Profiler 960 (EnzyScreen BV, Heemstede, The Netherlands). Frozen glycerol stocks were inoculated in 100 mL YPD medium and grown overnight. 0.5 mL of the overnight culture was transferred to 100 mL SMD + Ura/Lys/Ade and grown until the OD_660_ had doubled at least once to ensure exponential growth. From this culture a 96-wells microtiter plate (EnzyScreen, type CR1496dl) containing either SMD + Ura/Lys/Ade or SMEG + Ura/Lys/Ade with final working volumes of 250 μL was inoculated with a starting OD_660_ of 0.1. Growth rate analysis and data analysis was performed as described in Boonekamp *et al*. (37).

#### Respiration assay

Specific rates of oxygen consumption were measured in 4 mL volume in a stirred chamber at 30 °C. The oxygen concentration was measured with a Clark-type oxygen electrode (YSI, Yellow Springs, OH). Strains were grown overnight in SMD + Lys/Ade. The overnight culture was diluted twice in 100 mL SMD + Lys/Ade and SMEG + Lys/Ade. Glucose cultures were grown for another three hours to ensure exponential growth, ethanol cultures were grown for another 48 hours to ensure that strains were fully respiratory.

Strains were spun down and washed twice to prevent carry-over of carbon-containing medium and resuspended in SM. For respiration assays on glucose, 25 OD_660_ units of culture were inoculated in a total volume of 4 mL in fully aerated SM + Lys/Ade to a final OD_660_ of 6,25. Air flow in the chamber was stopped and 20 μL of a 50 % (w/v) glucose solution were added to a final concentration of 14 mM. For respiration assays on ethanol 40 OD_660_ units of culture were inoculated in a total volume of 4 mL in fully aerated SM + Lys/Ade to a final OD_660_ of 10. Air flow in the chamber was stopped and 50 μL absolute

Ethanol solution were added to a final concentration of 210 mM. The percentage of dissolved oxygen was measured until oxygen was depleted and was used to calculate the rate of oxygen consumption expressed as the decrease of oxygen in the reaction vessel over time, divided by the biomass concentration in the vessel.

#### DNA Sequencing and bioinformatics

Whole genome sequencing of strains 161-U7 and 161-I^0^ was performed by Macrogen Europe (Amsterdam, the Netherlands). Genomic DNA of samples 161-U7 and 161-I^0^ were sequenced at Macrogen on a Novaseq 6000 sequencer (Illumina, San Diego, CA) to obtain 151 cycle paired-end libraries with an insert-size of 550 bp using TruSeq Nano DNA library preparation, yielding 2 Gigabases in total per sample. Reads of both samples were mapped using BWA (version 0.7.15) (57) against CEN.PK113-7D genome. Alignments were processed using SAMtools (version 1.3.1) (51). *De novo* assembly was performed using SPAdes (version 3.9.0) (58) for both samples. Assembled contigs were aligned with nucmer (MUMmer package v3.1) (59) to CEN.PK113-7D reference, the overlapping contigs aligned to the mitochondrial chromosome were merged into scaffolds to reconstruct the mtDNA for both samples.The contigs containing *COX1* and *COB* as well as the consensus sequences of the mitochondrial genomes of 161-U7 and 161-I^0^ were aligned to the CEN.PK113-7D genome using BLAST (NCBI, USA) and visualized in R using genoPlotR (file: *alignments*.*Rmd*) (60). The sequencing data and *de novo* assemblies are deposited at NCBI (https://www.ncbi.nlm.nih.gov/) under BioProject ID PRJNA902953.

## Results

### 1. Isolation of mitochondria and preparation of RNA

As compared to genomic DNA-encoded transcripts, the quantification of all RNA species in mitochondria presents specific technical challenges that require tailor-made experimental and computational procedures. Firstly, the cellular abundance of mitochondrial RNA in yeast is very low and only represents circa 5 % of the total cellular RNA pool under respiratory conditions, and approximately 1 – 3 % under glucose repression, of which 90 – 95 % represents mitochondrial rRNA (21,61). Secondly, the absence of polyadenylation of yeast mitochondrial RNA prevents their enrichment by standard, polyA-tail based methods. Finally, to fully capture the diversity in mitochondrial RNA splicing variants, processing of the RNA should be kept to a bare minimum, as PCR or cDNA synthesis reactions might introduce biases. To overcome these problems, a mitochondrial RNAseq workflow was developed and tested (Figure 2A). Enrichment for mitochondrial RNA was achieved by physically separating mitochondria from yeast cells and extracting RNA from isolated mitochondria. To minimize steps that might bias the mitochondrial RNA pool, either by enriching for specific species or by modifying transcripts length, Nanopore long read direct RNA-sequencing was used, in which a single (polyadenylation) step is required between RNA extraction and sequencing. Synthetic control RNAs were spiked at three different stages to monitor mitochondrial RNA quality during extraction, polyadenylation, and sequencing (Figure 2A).

Yeasts cultures were spheroplasted and homogenized, and mitochondria were isolated from the other cellular fractions by differential centrifugation. Oxygen uptake measurements and microscopic analysis confirmed that isolated mitochondria were abundant and intact (Figure S 1). Western blotting and enzyme assays further showed that mitochondria were significantly enriched, but not fully devoid of cytosolic contamination (Figure S 1). The mitochondrial fraction can be contaminated by carry-over of cytosolic RNA species suspended in the cytosol and by RNA species attached to the outer mitochondrial membrane. These contaminants could be removed by RNAse treatment and by removal of the mitochondria outer membrane (i.e. mitoplasting). These treatments did however not significantly reduce the contamination by most cytosolic RNA and reduced mitochondrial RNA yields (Figure S 2). RNA was therefore directly extracted from crude isolated mitochondria without additional processing.

To quantify the contamination by non-mitochondrial RNA species and monitor RNA quality during extraction, two synthetic non-polyadenylated control RNAs were spiked to mitochondria prior to RNA extraction. A short (60 bp) *in vitro*-synthetized control simulated a tRNA-length transcript (IVT01), while the 1 kb eYFP transcript simulated messenger RNAs (IVT-eYFP, Figure 2A). Control RNAs of various lengths were also spiked further down the workflow. Spiking before poly(A)-tailing assessed the IVT reaction yield and the poly(A)-tail length, and a control supplied by Oxford Nanopore Technologies (ONT) was added prior to sequencing to assess sequencing throughput, and library size and quality (Figure 2A). Polyadenylation was required to make the mitochondrial and *in vitro* synthesized RNA compatible with the Direct RNA sequencing protocol of ONT, as the adapters required for Nanopore sequencing are added through a 10(dT) primer sequence that hybridizes with the poly(A)-tail of mRNA. As mitochondrial RNA is not natively polyadenylated, addition of a poly(A)-tail of at least ten adenine residues was required for successful adapter ligation and subsequent sequencing the RNA.

Following the above-described protocol, RNA-seq was performed using independent triplicate cultures of the *S. cerevisiae* laboratory strain CEN.PK113-7D grown with glucose or a mix of ethanol and glycerol (further referred to as ethanol media) as sole carbon source. RNA sequencing resulted in library sizes ranging between 0.4 to 1.1 Gb of data, with 0.4·10^6^ to 1.2·10^6^ reads that passed a quality score of above 7 (Spreadsheet S1). The average read N50 score was around 1.2 kb, indicating that most of the reads covered full gene length. The longest reads were around 8 kb, revealing that some, but not all full-length polycistronic primary transcripts were detected (expected size 1.6 to 16 Kbp, Figure 1). ONT Direct RNA sequencing does sequence poly(A)-tails of the RNA molecules, thereby enabling the quantification of poly(A)-tail length. The enzymatically added poly(A)-tails of mitochondrial and control RNAs were on average 40 - 50 adenine residues long, a length comparable to cytosolic RNA species (Figure S 3). 95 % of the basecalled reads passed the ONT quality control (Q-value >7) and all passed reads mapped to the reference genome. The mitochondrial RNA species represented around 5 - 12 % of the total number of passed reads, excluding the fraction of control reads, this represented a *ca*. twofold enrichment of mitochondrial RNA in both conditions, the rest being mostly cytosolic ribosomal RNA (Figure 2B,C, Figure S 4). The libraries were nevertheless large enough to comfortably cover the mitochondrial genome: the obtained datasets had on average 5·10^4^ reads that mapped to the mitochondrial genome, equating approximately 600 Mb of sequencing data to cover the 86 kb mitochondrial genome. On ethanol, the breadth of coverage was at an average of 75 % of the heavy strand and 15 % of the light strand of the mitochondrial genome, and all the open reading frames (ORFs) were covered by sequencing reads. Direct RNA sequencing therefore yielded enough reads of quality and of the expected size and was further explored to characterize the mitochondrial transcriptome.

### 2. Read distribution and length

A custom bioinformatics pipeline was set up to analyze the basecalled reads. In a first step, reads were aligned to the fully annotated CEN.PK113-7D nuclear and mitochondrial reference genomes (Figure 2). All expected RNA types from both mitochondrial and cytosolic origin were identified: mRNA, tRNA, rRNA, and noncoding RNA (ncRNA) which includes the RNA component of RNaseP *RPM1* and RNA sequences required for RNA-primed mitochondrial origins of replication (*ori*). Transcripts were categorized by type and ORF based on their mapped location on the genome, allowing the quantification and analysis of the origin and type of each sequenced transcript (Figure 2B). Most mitochondrial reads (90 – 94 %) belonged to rRNA species. All expected rRNA, mRNA and ncRNA were detected, although some in low abundance.

The mitochondrial transcriptome is characterized by a large diversity in length, the shortest being a 71 bp tRNA and the longest a 16.5 kb polycistronic transcript. The length and sequence of transcripts varies depending on processing of polycistronic transcripts and splicing of the respective genes. With the exception of very short RNAs, long-read RNAseq has the capability to capture this mosaic RNA landscape. For most mRNAs, the average read length was within 75 – 125 % of the expected ORF length, meaning that mRNAs were mostly sequenced in a single read (Figure 3A, Figure S 5, Figure S 6). Slightly shorter read length might reflect partial degradation of the RNA, while longer reads can reflect incomplete RNA processing or the presence of untranslated leader regions on the RNA. A few reads were substantially shorter or longer than the expected ORF length. Notably, the read length for the *ATP8* and *OLI1* genes was respectively six and three times longer than the expected mRNA length. (Figure 3A). In line with earlier reports, closer inspection of the read alignment of these loci confirmed the identification of a bicistronic transcript encoding *ATP6* and *ATP8* (62). However, long *ATP8* transcripts were also due to the presence of 5’ untranslated leader (UTL) and 3’ untranslated region (UTR) sequences, rather than polycistronic expression with the surrounding genes: 44 % of the reads mapping to *ATP8* were bicistronic, whereas 55 % of the reads were extended due to attached UTL and UTR regions (Figure 3D). *OLI1* is part of a 4.8 kb polycistron, however while a few transcripts extending past the *OLI1* dodecamer were detected (Figure 3E), no full-length primary transcripts were found. In agreement with Turk *et al*., *OLI1* transcripts elongation was caused by the presence of a 5’UTL (13).

**Figure 3.**
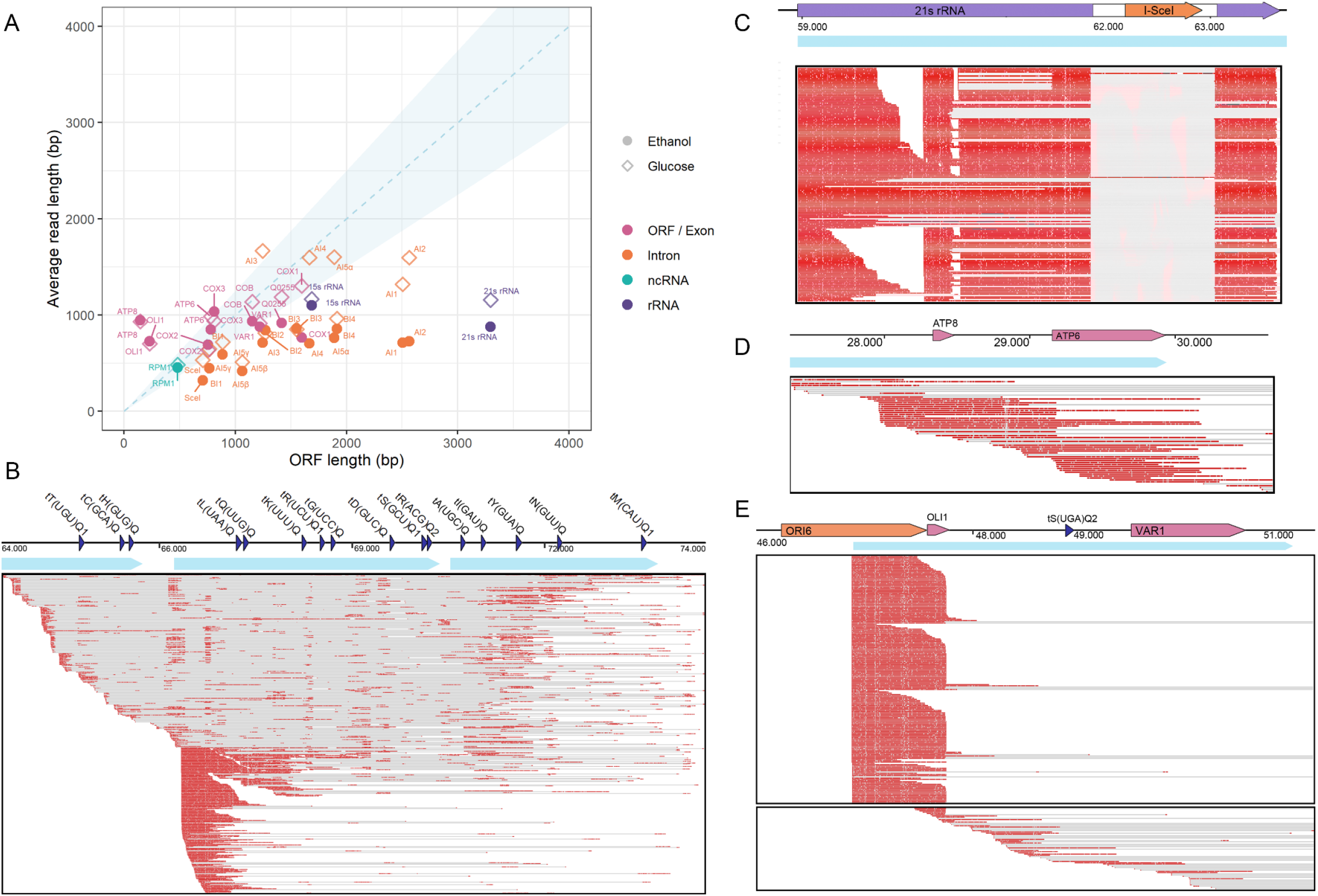
Read length and coverage of genes. A) Average read length for each gene plotted versus the expected ORF length, where the ORF length is defined as the length of a matured RNA from start- to stop codon, after splicing (if applicable). Reads from ethanol grown cultures are shown as closed circles, reads from glucose-grown cultures as open diamonds. Reads are separated per type, mRNA: messenger RNA (pink), ncRNA: non-coding RNA (green) rRNA: ribosomal RNA (purple). Spliced genes are separated between the main exon (pink) and the different spliced-out introns (orange). The dashed line represents a full-length read (gene length = read length), the blue shaded area represents an interval where the read length is 75 % to 125 % of the full gene length. B, C, D, E) Visualization of raw reads of one representative replicate on ethanol, aligned using Tablet with the primary transcripts of the tRNA cluster (B), 21s rRNA (C), ATP8-6 (D) and OLI1-VAR1 (E). The colored arrows indicate the genes and their respective locations, the primary polycistronic transcripts are indicated with light blue arrows. Boxes show visualization with Tablet of sequencing reads mapped to the different loci, a red line indicates correct mapping of a sequencing read to the location on the mtDNA, a grey line indicates a gap in the read.

Non-coding RNA sequences were in general shorter than their respective ORF length. The most extreme case was 21s rRNA for which less than 1 % of the reads mapping to 21s rRNA gene displayed the expected gene length. Since the used ORF lengths already exclude intron sequences, this cannot be the result of splicing. Analysis of the raw reads mapped to the 21s rRNA locus revealed that reads were strongly truncated or degraded at approximately 1 – 1.5 kb after the 5’ start of the 21s rRNA gene (Figure 3C). Additionally, most of the intron sequences of *COX1* and *COB* (*aI1-aI5γ* and *bI1-4* respectively) were shorter than their expected length. This can be explained by the rapid turnover of spliced introns, essential for efficient protein expression and respiration and to prevent intron accumulation that may cause intron-toxicity by catalyzing ‘exon-reopening’ reactions (63,64).

The fraction of tRNA’s in the present dataset was particularly low (Figure 2B). Processed tRNAs have a short length, ranging from 71 to 90 bp. The data showed that the spiked RNA controls below 102 bp were strongly depleted during the RNA-seq protocol (Figure S 7). This result was in line with earlier reports that Nanopore sequencing cannot accurately analyze short nucleotide sequences of < 100 bp (65) and suggests that short RNAs such as tRNAs were underrepresented in the present study. All tRNA’s are part of a polycistronic transcript, either in a tRNA cluster or grouped with other RNA transcripts.

Analysis of reads aligned to the tRNA cluster revealed that single processed tRNA’s were indeed rarely detected and that any captured tRNA’s were part of a polycistronic transcript (Figure 3B).

Based on the nonanucleotide sequences that initiate translation in the mitochondria, it is assumed that the mitochondrial genome is expressed in 11 primary polycistronic transcripts that are further processed at internal dodecamer sites and by endolytic cleavage of tRNA’s. None of these full length polycistronic transcripts were detected, the longest read was 8 kb and mapped to the *COX1* pre-mRNA. However, knowledge on processing of these polycistronic transcripts is still very limited, and polycistronic transcription has only been demonstrated indirectly (62,66). It was only recently found that the dodecamer is recognized by the protein Rmd9p and that deletion of this protein results in detection of unprocessed primary transcripts (67,68). Also, processing may be co-transcriptional and therefore difficult to capture *in vivo*. Nevertheless, the read length analysis revealed some discrepancies between read length and gene length that suggest polycistronic expression of the mitochondrial genome.

### 3. mtRNA coverage and landscape

The present datasets originated from three biological culture replicates, mitochondria isolation rounds and sequencing runs, leading to different read distributions and library sizes (Figure 2B,Table S 5). Quantitative analysis of expression levels therefore required library normalization. RPKM (Reads Per Kilobase Million) is the standard normalization method for RNA-seq, developed for short-read RNA sequencing in which read length (*ca*. 150 - 200 bp) is substantially shorter than gene length (e.g. several Kb in yeast). RPKM normalizes to library size, but also to gene length, as to compensate for this direct correlation between read number and gene length. However, normalization by gene length is not applicable to long-read RNA sequencing as read length often approaches or even exceeds the gene length. The datasets were therefore only normalized over library depth in Counts-per-Million (CPM).

The sum of polycistronic primary transcripts of the mtDNA encompassed 54.1 Kbp of the heavy strand, which means that 64 % of the heavy strand is theoretically expressed, and 6.3 kB (7 %) of the light strand. In the obtained RNAseq dataset, the breadth of coverage of the mitochondrial DNA heavy strand was higher than expected, with 74.7 % and 85.7 % for ethanol- and glucose-grown cells respectively (Figure 4A,B, Figure S 8, Figure S 9), revealing a very good representation of the mitochondrial transcriptome When looking at the raw read alignment of coding and non-coding regions, the respective 10 – 20 % higher expression does not have a clear origin and seems to mostly result from a combination of occasional read-through past the dodecamer or mis-alignment of read ends (Figure S 10), which was especially prevalent in the AT-rich 3’end of 15s rRNA (Figure S 11).

**Figure 4.**
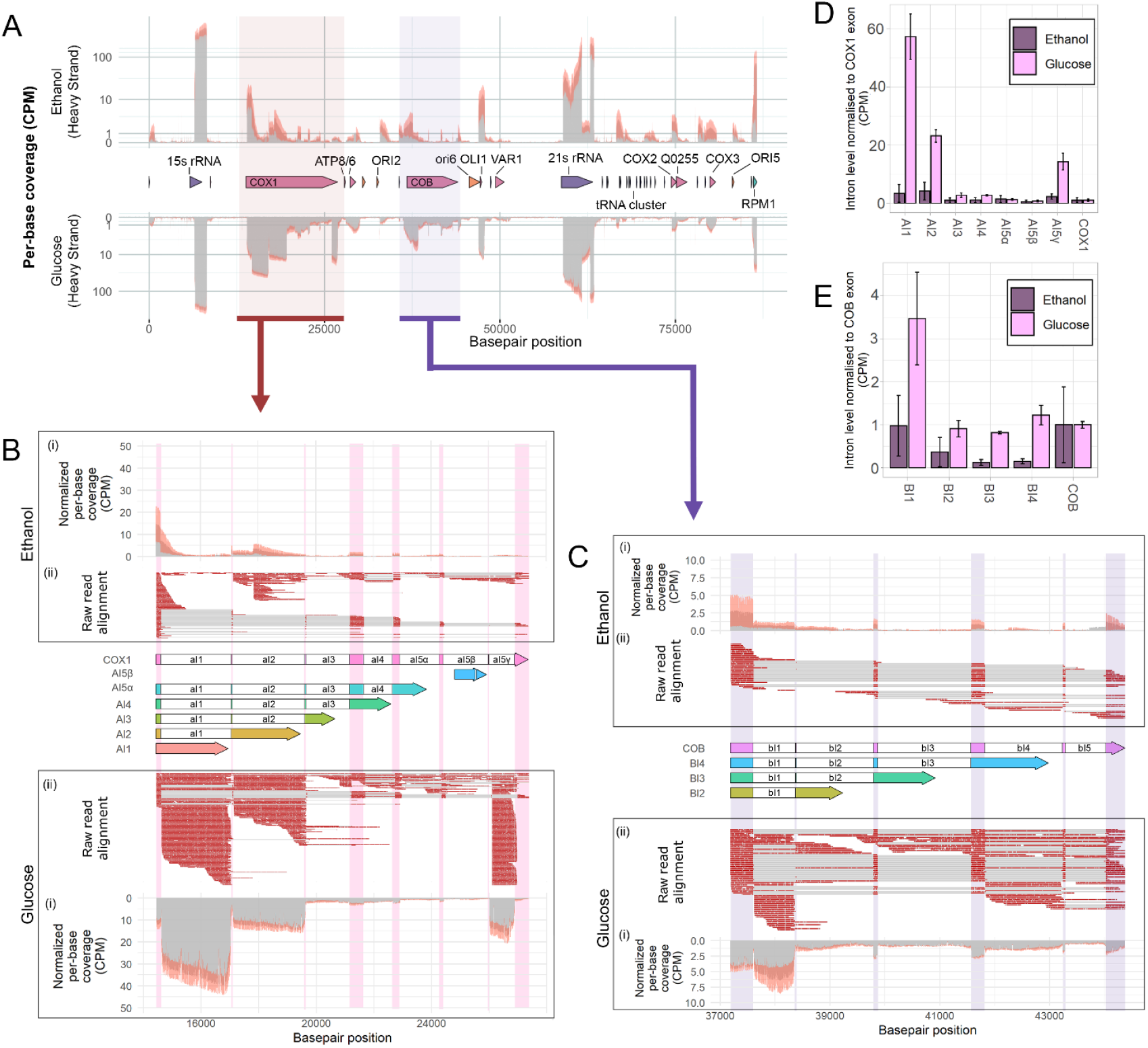
Sequencing coverage of the mitochondrial transcriptome of cultures grown on ethanol and glucose. A) per-base coverage of the mitochondrial transcriptome of cultures grown on ethanol (top) compared to the per-base coverage of the mitochondrial transcriptome of cultures grown on glucose (bottom), normalized per million bases as Counts-Per-Million (CPM) of the heavy (forward) strand of the mitochondrial DNA. Coverage depth is represented in grey, the standard deviation between the sequencing depth of triplicate experiments is shown in orange. B) schematic overview, sequencing depth and splicing of the COX1 Open reading frame (ORF). The middle shows the COX1 ORF, where the exon sequences are shaded in pink, colorless areas are spliced out of the final COX1 mRNA. The different splicing variants of the COX1 open reading frame are shown below the COX1 ORF. Capitalized names are used to indicate protein-encoding introns, non-coding introns are shown in lowercase. (i) shows per-base coverage of the COX1 ORF, normalized per million bases as Counts-Per-Million (CPM). Coverage depth is represented in grey, the standard deviation between the sequencing depth of triplicate experiments is shown in red. (ii) shows visualization of sequencing reads mapped to the COX1 ORF visualized using Tablet, a red line indicates alignment of a sequencing read to the location on the ORF, a grey line indicates a gap in the read. The analysis was done for mitochondria of ethanol-grown cultures (top) and glucose-grown cultures (bottom). The location of the COX1 exon is shaded in pink for visualization purposes. C) schematic overview, sequencing depth and splicing of the COB ORF. D) Transcript levels of the COX1 introns on glucose (light pink) and ethanol (purple), normalized to library size in CPM and normalized to transcript levels of the COX1 exon mRNA. E) Transcript levels of the COB introns on glucose (light pink) and ethanol (purple), normalized to library size in CPM and normalized to transcript levels of the COB exon mRNA.

For the light DNA strand, only containing one active origin of replication and tRNA tT(UAG)Q2, the observed coverage was 14.8 % for ethanol-grown and 7 % for glucose-grown cultures (Figure 4A, Figure S 8, Figure S 9). Most expression clearly originated from the expected primary transcripts. On ethanol, the breadth of coverage of the light strand was slightly higher than expected, but the depth of this coverage was very low. This might result from transcriptional read-through at the *ori3* locus or from the presence of mirror RNA’s, which have been described to exist in similar low quantities in both human and yeast mitochondria and are possibly a by-product of RNA processing (13,69).

On ethanol, all of the coding regions were sufficiently covered by sequencing reads (Figure 4, Figure S 10), including regions encoding tRNA’s, albeit in a low amount and often as polycistronic reads (Figure 3B). The gaps in coverage represented 14 – 25 % of the mitochondrial genome, but all in non-coding regions. The mitochondrial genome was therefore exhaustively sequenced, despite the low representation of mitochondrial RNAs in the total RNA-seq data (*ca*. 7%, Figure 2B). Irrespective of the carbon source used, the per-base coverage showed a large variation in expression between the mitochondrial genes (up to 3300-fold difference, Figure 4A). The described method yielded a complete coverage of the mitochondrial genome and could therefore be used to quantify changes in RNA expression and processing of the mitochondrial genome between different conditions.

### 4. Comparison of transcript levels between ethanol and glucose-grown cultures

While comparing the relative expression of the various mitochondrial RNA species within a given growth condition is straightforward and only requires normalization over library depth in Counts-per-Million (CPM), comparing RNA abundance between growth conditions is more challenging. *S. cerevisiae* is a typical Crabtree positive yeast, in which excess glucose triggers a respiro-fermentative metabolism. Glucose exerts a strong repression on respiration, leading to low respiration rates, small and sparse mitochondria, and lower mitochondrial RNA levels (21,70,71). Conversely, ethanol is fully respired, leading to abundant and very active mitochondria. The present method first extracted mitochondria from different culture volumes on glucose and ethanol, while RNA sequencing is performed on the same input amount of RNA. Comparing mtRNA levels during growth on glucose and ethanol therefore requires an additional correction for the abundance of mitochondrial mass per cell, between the two carbon sources. Since direct RNAseq results in smaller library sizes (∼ 0.5-1M aligned reads) well below the required input to obtain a high-power differential gene expression using short-read sequencing (∼ 10 M reads), gene expression was measured as transcript-level counts (18,72,73). Additionally, a recent study reported a correlation between mitochondrial mass and cell dry weight of glucose-grown and ethanol-grown cultures (43). This correlation was used to normalize the counts per mitochondrial dry weight, thereby enabling the direct comparison of mtRNA abundance between glucose- and ethanol-grown cultures (Figure 5, Table S5).

**Figure 5.**
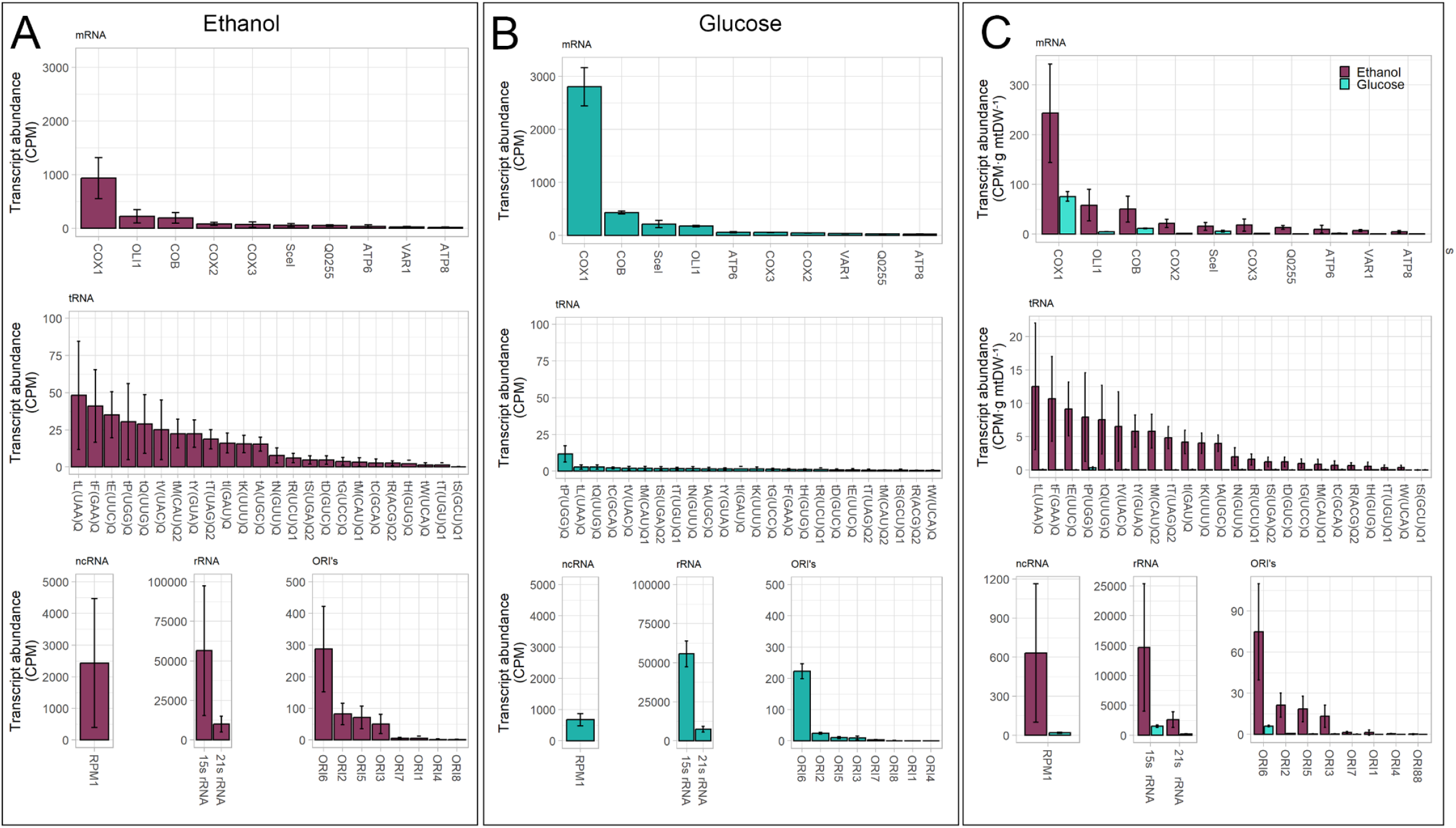
Transcript abundance of mitochondrial RNA species normalized to library depth in Counts-per-Million (CPM) of mitochondria isolated from ethanol-grown cultures (A) and glucose-grown cultures (B). Error bars indicate the standard error between triplicate experiments. Reads are grouped per type, mRNA: messenger RNA (no separation made between introns and exon reads), ncRNA: non-coding RNA, rRNA: ribosomal RNA, ori: replication origin. C) Comparison of transcript abundance of mitochondrial RNA species of mitochondria isolated from ethanol-grown cultures (dark purple) and glucose-grown cultures (light blue), normalized to library depth in Counts-per-Million (CPM) and normalized to grams mitochondrial dry weight of the cultures (g mtDW).

In both conditions, based on non-normalized data, the most abundant RNA species were rRNAs, the 15s rRNA being 5.5 times more abundant than the 21s rRNA (Figure 5A, B). Transcripts of all mitochondrial mRNAs were recovered in both conditions, but their relative expression was affected by the type of carbon source. For instance, transcripts for the cytochrome *c* oxidase subunits 1, 2 and 3 (*COX1, COX2* and *COX3*, respectively), the cytochrome *b* (*COB*) and the F0-ATPase subunit c (*OLI1*) were most abundant in ethanol-grown cultures (Figure 5A). Conversely, on glucose *COX1* was the most abundant mRNA, followed by *COB* and the meganuclease-enconding transcript I-SceI, encoded in the omega-intron of 21s rRNA (Figure 5B).

In line with the reduced involvement of mitochondria in respiration with excess glucose, the overall normalized abundance of mtRNA was lower in glucose-than in ethanol-grown cultures. Accordingly, with the notable exception of *COX1* and *COB*, the abundance of transcripts encoding respiratory chain subunits was 5 to 15 times higher in ethanol-than in glucose-grown cultures (Figure 5C, Figure S 12). The high expression of *COX1* relative to *COX2* in glucose conditions was unexpected. *COX1, 2* and *3* together encode the catalytic core of cytochrome *c* oxidase and exist in a 1:1:1 stoichiometry in the inner mitochondrial membrane (74-76). The presented finding also contradicts a previous study on the stoichiometry of mitochondrial protein- and RNA levels under glucose repression (21,43).

Additionally, tRNAs were more abundant in ethanol-than in glucose-grown cultures, which is in line with the increase in protein synthesis in ethanol-grown cultures and previous studies on tRNA levels under glucose repression (77). Out of the 24 tRNAs, 22 could be quantified in all ethanol-grown cultures, while only tP(UGG)Q was reliably detected in glucose-grown cultures (Figure 5). In line with these observations, the third most abundant RNA species in ethanol-grown cultures was *RPM1. RPM1* encodes the RNA component of RNase P, an enzyme highly active during respiratory growth that is essential for tRNA processing (78,79). All tRNA’s are part of a polycistronic transcript, either in a tRNA cluster or grouped with other RNA’s, and the three most abundant RNA species tL(UAA)Q, tF(GAA)Q and tE(UUC) are located at the start of a polycistronic primary transcript (Figure 1).

The dataset also provides insight on the expression of the origins of replication. It is still unclear whether mitochondrial DNA replication relies on RNA priming, rolling-circle replication, or a combination of the two (80,81). Nevertheless, transcripts for three (*ori*2, *ori*3, *ori*5) out of the eight mitochondrial origins of replication were detected under both growth conditions, suggesting that these origins were active under the tested conditions (Figure 4A, Figure 5). This is accordance with the presence of uninterrupted promoter sequences for these *ori*’s (13,82-84). The abundance of these replication primers was three- to five-fold higher with ethanol as carbon source (Figure 5). Many reads also mapped to *ori6*, which was most likely an artefact caused by the presence of the *OLI1*/*VAR1* primary transcript within this origin. This hypothesis was supported by the lack of reads mapping to the 5’-end of ori6 (Figure 3E).

Remarkably, the read coverage within genes was also carbon-source dependent, particularly for the *COX1* and *COB*, and 21s rRNA loci (Figure 4A). For instance, the coverage within the 21s rRNA on ethanol as compared to glucose shows a sudden decrease in between the 5’ end and the omega-intron, at approximately 1 Kb after the 5’ end, which corresponds to the previously observed gap in read alignment (Figure 3C). This gap suggests that reads might be truncated or degraded from this point onward. The fact that the dip in coverage originated from the same position suggests that shearing of RNA during processing is not a likely cause, as this RNA damage would occur randomly. A similar dip in coverage was observed for the *RPM1* gene, for which the decrease in coverage and read truncation exactly matched the RNA processing pattern and proposed heptakaidecamer cleavage site (Figure S 13, (13)). Taking this into consideration, the coverage pattern observed for 21s rRNA might suggest the presence of an RNA processing site.

Long read sequencing can therefore be used for the quantitative comparison of transcript levels between growth conditions. However, the polycistronic nature of the long RNA reads requires careful interpretation of the data taking into consideration the mapped location and coverage within a gene of the reads.

### 5. Changes in splicing patterns of *COX1* and *COB* between growth on ethanol and glucose

Comparison of the read coverage on glucose- and ethanol-grown cultures showed pronounced differences for *COX1* and *COB* (Figure 4A-C), with a 5 to 50-fold increase in coverage of sub-sections of these ORF’s when glucose was the carbon source. A uniform coverage over the full ORF is not expected, since the *COX1*- and *COB* ORF’s do not only encode subunits of cytochrome *c* oxidase and cytochrome *b*, respectively but also contain several introns and intron-encoded genes (Figure 4B,C). The *COX1*-ORF (or pre-mRNA) is 12.9 Kb long, of which 1.6 Kb consist of exon sequences that together encode the Cox1p. The *COB* pre-mRNA is 7.1 kB long of which 1.2 kB encode Cobp. The *COX1* and *COB* introns belong to either group I or group II homing introns that are distinguished by their splicing mechanism and are considered phenotypically nonessential for *S. cerevisiae* (16,85,86). The *COX1* exon sequences are interrupted by four group I and three group II intron sequences (*COX1-aI1, aI2, aI3, aI4, aI5α,β,γ)*, and *COB* contains one group I and four group II intron sequences (*COB-bI1, bI2, bI3, bI4, bI5)*, that are spliced out of the pre-mRNA in a protein-facilitated manner (16,31,87). As a result of alternative splicing (part of) these introns can also encode proteins (Figure 4B,C). *COB*-*BI2*, -*BI3* and –*BI4* encode maturases, *COX1*-*AI1* and –*AI2* encode reverse transcriptases, and *COX1*-*AI3*, -*AI4* and -*AI5α* encode endonucleases, all of which facilitate splicing and intron mobility of their respective intron sequence (8,16,87,88). Finally *COX1-AI5β* encodes a putative protein of unknown function, group II introns *aI5γ* and *bI1* are non-coding.

In ethanol-grown cultures, the pre-mRNAs of both *COX1* and *COB* were clearly spliced and processed to yield an RNA containing only exon sequences (Figure 4B,C). For *COX1*, spliced intron sequences were also detected, but displayed various degrees of degradation and were rarely found in full-length, in agreement with the shorter than expected read length of intron sequences (Figure 3A). Overall, no significant difference was observed between intron and exon abundance of *COX1* in ethanol-grown cultures (Figure 4B,D). For *COB*, intron sequences were rare, especially introns *bI3* and *bI4* were strongly depleted and present at only 10 % of the exon level. Some reads showed patterns of alternative splicing, but since intron levels are relatively low and intron turnover is co-translational in mitochondria (89), it is difficult to attribute the intron levels to either ‘regular’ (*aI1-5, bI1-5*) or alternative splicing (*AI1-5, BI2-4*) of the intron.

In glucose-grown cultures, the splicing pattern of the mature *COX1* and *COB* exon was more difficult to infer from the raw read alignment, and unspliced pre-mRNAs were clearly more abundant (Figure 4B,C). Also in contrast with ethanol-grown cultures, in glucose-grown cultures the coverage of several sequences mapping to intronic regions was substantially higher than the corresponding exon abundance for both *COX1* and *COB* (Figure 4B,C). *COX1*-*aI1*, -*aI2* and -*aI5γ* were respectively 57-, 23- and 14-times more abundant than the mature *COX1* exon-RNA (Figure 4D). Similarly, the *COB-bI1* intron was 3.5-times more abundant than the *COB* mature exon-RNA (Figure 4E). Di Bartolomeo *et al*. (43) reported that the abundance of the Cox1 protein was 15 to 50 times higher than its intron-encoded proteins, a result in contrast with the relative exon-intron abundance measured in the present study (Figure S 14). Additionally, analysis of the raw read alignment to the *COX1* and *COB* ORFs revealed that the increased intron levels were not due to alternative splicing, as the intron sequences were rarely attached to a 5’ exon sequence (Figure 4B,C). Lastly, considering that the *aI5γ* and *bI1* introns do not encode any protein (16), the high intron abundance most likely results from the accumulation of the spliced intron sequences rather than the elevated expression of intron-encoded genes.

Strikingly, all of the introns with increased levels on glucose (*COX1-aI1,2,5γ, COB-bI1*) belong to the group II of introns, while the levels of group I intron sequences (*COX1-aI3,4,5α,5β* and *COB-bI2,3,4*) remained similar to the level of exon RNA (Figure 4D,E). Group II introns are circular RNAs (known as lariats) and are a result from the covalent linking between the 5’ end and an A/U-residue 8 to 9 base pairs from the intron 3’ end. The formation of this branch point is followed by exon ligation and cleavage at the 3’ splice site and the resulting release of a circular intron lariat (16,90,91). Nanopore technology cannot sequence circular RNA molecules, which means that (part of) the lariats were most likely linearized prior to sequencing, either *in vivo* or during sample processing. Introns *aI1* and *aI2* are 2.3 Kb in length, while *aI5γ* and *bI1* are both around 0.8 Kb. Over 75 % of reads for the *COX1-aI1, -aI2, aI5γ* and *bI1* intron had the expected length up to the branch point, which indicates little to no degradation (Figure 4B,C). Additionally, the median weighted read length (N50) of the dataset was 1.4 Kb and most of the RNA was sequenced in full (Figure 3A), meaning RNA degradation during library preparation was likely limited. Lastly, mechanic shearing of the circular RNA during library preparation would result in a random location of the break for each RNA molecule. When comparing the alignment of the raw reads to each intron, no random breakage of reads was observed, as reads either consistently covered the full intron from 5’ to 3’ end or were partially degraded from the 5’ end (Figure 4B,C).

This data suggested the accumulated group II introns of mitochondria of glucose-grown cells were not sheared and were sequenced in full, up to the branching point. Approximately half of the reads for these introns included the branch point sequence, with only very few (5 to 10 reads per sample) surpassing the branch point (Figure S 15, Figure S 16, Figure S 17, Figure S 18). Additionally, the abundance of the group II intron sequences is likely underrepresented in both glucose- and ethanol-grown cultures, as circular lariat molecules were most probably present in the isolated RNA, but were not available for sequencing due to the nature of Nanopore technology.

### 6. Exploring the effect of the presence of group II introns on yeast physiology

Group II introns specifically accumulated during respiro-fermentative growth in glucose medium. While this accumulation might be a side effect of some yet unknown condition-dependent regulation of intron stability, the possibility of a physiological role for mitochondrial lariats cannot be excluded. In higher eukaryotes, there are indications that noncoding RNA can play a role in gene expression, epigenetic regulation, and genome stability (92-94). The hypothesis that intron sequences may play a similar role in yeast can be tested by comparing the physiology of yeast strains with and without group II introns. Despite the poor genetic accessibility of the mitochondrial genome, a few ‘intron-less’ strains have been previously constructed (31,95). The physiological response of an intron-less strain from the 161 strain lineage (also known as ID41-6/161) to growth in the presence of glucose or ethanol was therefore explored (29,30). The intron-less strain 161-I^0^ does not contain introns in *COX1* and *COB* and also lacks the ω-intron (encoding I-SceI) (31). The control congenic strain 161-U7 just lacks the ω-intron.

As the sequence of these strains’ genome was unavailable, *S. cerevisiae* 161-U7 and 161-I^0^ genomes were sequenced using Illumina short-read technology. A *de novo* assembly was performed based on the sequencing reads (Table S 6), and resulting contigs containing *COX1* and *COB* could be aligned to the CEN.PK113-7D mtDNA to determine the absence of intron sequences (Figure 6A). Additionally, a consensus sequence of the mitochondrial genomes of 161-U7 and 161-I^0^ was generated by mapping the contigs from the *de novo* assembly of the two strains to the CEN.PK113-7D mitochondrial genome and extracting the resulting consensus sequence (Figure S 19). In addition to the mitochondrial genome, the complete sequence of the nuclear genome of these strains was *de novo* assembled and is available at NCBI (see M&M section).

**Figure 6.**
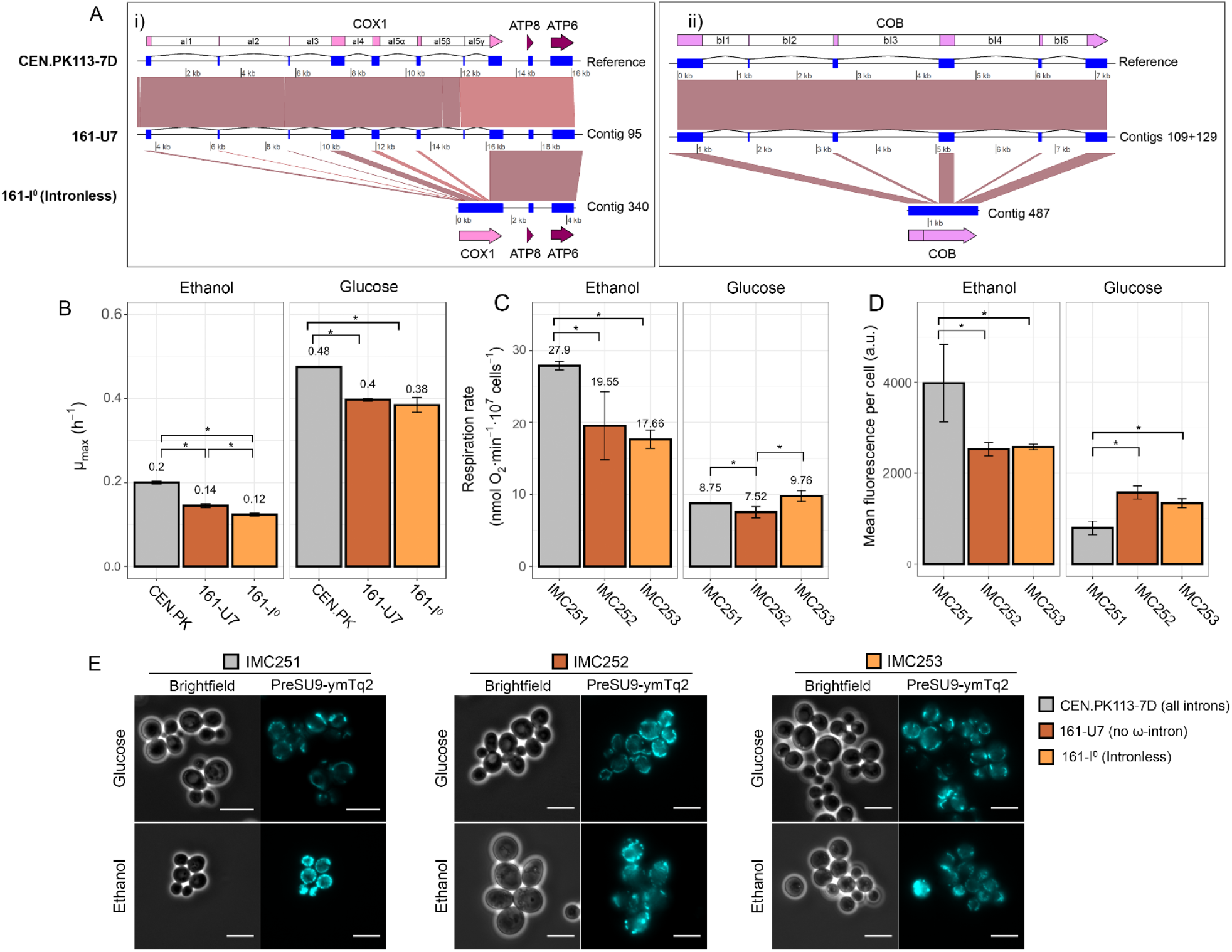
Comparison of the genomes and physiology of the CEN.PK113-7D, 161-U7 and 161-I^0^ strains. A) Alignment of the i) COX1 and ii) COB loci on the mitochondrial genome of CEN.PK113-7D, and contigs of 161-U7 and 161-I^0^. COB of 161-U7 was divided in 2 contigs which were combined at a 76-bp overlap. The mitochondrial genome is represented as a black line and the respective coordinates are indicated below each genome/contig. The locations of exon mRNA, rRNA and ncRNA are indicated as blue squares, tRNA and ori are not shown. Spliced exons are connected by a line, annotations of the sequences are indicated above (CEN.PK113-7D, 161-U7) and below (161-I^0^) the alignment. Grey boxes or lines indicate a sequence identity of > 95 % (BLAST) between the two alignments. B) Maximum specific growth rate (h^-1^) in microtiter plates of CEN.PK113-7D, 161-U7 and 161-I^0^ on ethanol and glucose medium. C) Respiration rate, D) Mean fluorescence intensity per cell measured in the mTurquoise2 channel, determined by microscopy and E) microscopy pictures of strains IMC251 (CEN.PK113-7D), IMC252 (161-U7) and IMC253 (161-I^0^) grown on SM-ethanol or SM-glucose with a blue fluorescent mTurquoise2 (mTq2) protein targeted to the mitochondria with a preSU9 mitochondrial targeting signal. Mean fluorescence was calculated per cell (n ≥ 250), the cell area was based on the brightfield image. Scale bars represent 10 μm. An asterisk (*) represents a significant difference with p< 0.05 (paired two-tailed student T-test).

Comparing the mitochondrial consensus sequences of the 161-U7 and 161-I^0^ confirmed the absence of the ω-intron in both strains (Figure S 19) and the absence of all other introns in strain 161-I^0^ (Figure 6). The mtDNA of 161-I^0^ covered 77 % of the 161-U7 genome and the similarity between the two mitochondrial consensus genomes was 99.2 % (Table S 7). The *de novo* assembly resulted in fragmented mitochondrial genomes (Table S 6) and must be complemented with long-read sequencing to confirm the correct structure of the mtDNA, therefore the analysis of the *COX1* and *COB* loci was performed using the correctly assembled *de novo* contigs that contained these loci rather than the generated consensus sequence (Figure 6). The similarity between the *COX1* and *COB* alleles was 99.8 and 99.9 % respectively (Table S 7), yielding 12 SNP’s in *COX1* and 1 in *COB*. Additionally, in the *COB* locus of intron-less strain 161-I^0^ the 14 bp exon sequence between introns *bI1* and *bI2* appeared to be removed from the mtDNA (Figure 6A).

A previous study reported substantial physiological differences between the two strains during growth on glucose, but growth on ethanol was not explored (96). In contrast with this earlier study, 161-U7 and its intron-less variant grew at similar rates on glucose medium. In cultures with ethanol as carbon source the intron-less strain grew slightly but significantly slower (14 %, p-value 0.05, student t-test homoscedastic) than its parent (Figure 6B). Such a change in growth rate may be explained by mutations that occurred in construction of strain 161-I^0^ or the absence of the second exon of *COB*. The respiration rate was similar for the two strains during growth on ethanol and was increased by *ca*. 30 % during growth on glucose for the intron-less strain (Figure 6C), which might indicate a partial alleviation of glucose repression. The two strains were equipped with a mitochondrial fluorescent protein to monitor mitochondrial mass. When applied to the CEN.PK113-7D control strain, this method showed a four-fold decrease in mitochondrial mass in glucose-grown as compared to ethanol-grown cultures (Figure 6D,E). This result was in good agreement with earlier reports for the same strain (43). The same method applied to 161-U7 and 161-I^0^ showed a substantially smaller mitochondrial mass variation between glucose- and ethanol-grown cells, where mitochondrial mass on glucose represented approx. 60 % of the mitochondrial mass on ethanol but revealed the absence of difference in mitochondrial mass between the two strains. The absence of introns did therefore not lead to clear physiological differences when comparing growth on glucose and ethanol media.

## Discussion

In the present study, RNA enrichment by mitochondria isolation combined with Nanopore long-read sequencing enabled the capture of the full mitochondrial transcriptome in the model eukaryote *S. cerevisiae*. All expected RNA species were found and the mitochondrial RNA coverage was sufficient to quantify most of the transcriptome. Small RNA molecules of below 100 bp, specifically tRNAs, were an exception, thus studies focusing on these RNA species should rather opt for, or complement with, short read sequencing, or use methods to specifically enrich these transcripts (97). The coverage of the mtRNA was sufficient, but can be further improved through removing contaminating RNA species by further purification of mitochondria (*e*.*g*. using Nycodenz gradient purification (98). Additionally, treatment with nucleases could reduce carry-over of cytosolic RNA, which represents up to 97 % of the sequenced RNA in the present study. Depletion of rRNA (99) is another approach to enrich samples for the relevant RNA species. Finally, the excessive number of reads originating from the RNA controls can be substantially reduced by decreasing the amount of spiked-in control RNA. Despite the polycistronic nature of the mitochondrial RNA, no full-length primary transcripts were detected. This result might reflect the real *in vivo* situation, in which the short half-life of polycistrons results from co-transcriptional splicing (89). However, although mitochondrial RNA is stabilized at its dodecamer sequences, it cannot be excluded that during sample processing of the intact mitochondria, the RNA processing and turnover enzymes stay active (19,67), particularly during spheroplasting which is performed at 30 &C for 30 – 45 minutes. Investigating the possibility of additional quenching or stabilization of the mitochondria prior to isolation may improve detection of unprocessed RNA (100,101).

While the present approach cannot give a perfect snapshot of *in vivo* RNA abundance, applying the exact same protocol to all samples quantified, for the first time, mtRNA abundance between different conditions, with good reproducibility between biological replicates (Figure 5). In a previous study, the transcriptome of *S. cerevisiae* grown in ethanol medium was monitored using short read sequencing (13). The relative abundance of RNA species from short-read sequencing is overall in good agreement with the present study. Small variations can likely be attributed to differences in RNA preparation methods, yeast strain genetic background, sequencing methods and normalization of the data (RKPM by (Turk *et al*.) and CPM in the present study). Notable exceptions were the expression levels for *15s rRNA* and *RPM1*, which were the first and third most abundant species in the presented study, with a 50- and two-fold higher expression compared to *COX1*, whereas Turk *et al*. reported the abundance of 15s rRNA to be comparable to *COX1* and *RPM1* approximately 10-fold lower compared to *COX1*. This may be explained by the secondary structure of these RNAs, which can interfere with binding of short sequencing reads, whereas Nanopore direct RNA sequencing includes relaxation of secondary structures (73,102).

As opposed to short-read sequencing, the present approach enabled the analysis and visualization of splicing variants with single base pair resolution, and the quantification of the abundance of RNA species at different processing stages. As transcripts were sequenced in a single read, RNA processing events could be identified by combining information on transcript coverage and on the presence of gaps in read mapping to the reference genome. This was best illustrated by comparing the known RNA cleavage sites of *RPM1* with the coverage and read alignment of that gene (Figure S 13). Similarly, the 21s rRNA showed a strong decrease in coverage and gap in read alignment in a single locus during growth on ethanol media. The consistent occurrence of the same phenomenon for all three independent replicates for ethanol-grown cultures, but absence for glucose-grown cultures, strongly suggested the existence of a condition-dependent new processing site for these RNA species. As the 21s rRNA sequence harbors no heptakaidecamer or internal dodecamer splicing sites, the exact target site for this novel RNA processing event remains to be defined.

The most remarkable difference between glucose- and ethanol-grown cultures was the substantially higher abundance (4- to 60-fold) of group II introns *COX1-aI1, -aI2, -aI5γ* and *COB-bI1* in glucose-grown cultures. Splicing of these lariat introns requires their circularization via internal covalent linking. However, Nanopore technology cannot sequence circular RNA molecules, and the 8-9 bp branch of the lariat is too short to be poly(A) tailed (90)^1^, which indicates that the fraction of introns that was sequenced exists in linear configuration. The existence of linear intron RNA in the mitochondria has been hypothesized but never demonstrated *in vivo* (77). Linearization can result from different mechanisms such as *in vivo* debranching by specific enzymes or degradation by endonucleases, or by shearing during sample processing. The reproducibility of group II introns abundance and length between biological replicates suggests that the difference in intron abundance observed between glucose- and ethanol-grown cultures was not an artefact caused by mechanical RNA shearing during sample processing, as approximately half of the reads covered the lariat branching point and were sequenced in full length (Figure S 15-Figure S 18), The only known intron-debranching enzyme in *S. cerevisiae* (Dbr1p (103)) is not localized to mitochondria (prediction using DeepLoc2.0, Table S 8 (104)) and Dbr1p has not been detected in any mitochondrial proteome (43,105). Therefore, it can be reasonably assumed that lariats are either debranched *in vivo* by a yet unidentified debranching enzyme, or that (a subset of) the group II introns is spliced as linear RNA molecules when grown on glucose. This may also mean that the presented intron levels are a underrepresentation of the actual intron level, as the introns with lariat structure could not be sequenced using Nanopore sequencing. Using *in vitro* debranching with purified Dbr1 could provide more insight on this.

Regardless of the configuration of the group II introns, a clear condition-dependent change in group II introns abundance was observed when cultures were grown on glucose. The most likely mechanisms leading to this change in intron-abundance are an increase in either intron stabilization by proteins on glucose or *in vivo* degradation of introns on ethanol. Potential target proteins may be identified by comparing the differential expression of RNA binding proteins observed between glucose- and ethanol-grown cultures (43,106,107). From this comparison, *YBR238C*, mitochondrial paralog of Rmd9p that stabilizes mitochondrial RNA at their dodecamer sequence (67,68), was 1.8-times more abundant in glucose-than ethanol-grown cultures (Figure S 14). MtRNA binding protein levels between ethanol and glucose were also compared relative to Cox1p expression, as this gives insight in the different levels of protein required between ethanol and glucose to mature a similar level of *COX1* mRNA. When normalizing to Cox1p levels, Mss116p, an RNA-helicase and –chaperone involved in splicing group II introns (31), showed over twofold increase in expression between growth on ethanol and glucose (Figure S 14), as well as Dis3p, which was demonstrated to play a role in mitochondrial intron decay (13). Combined with the difference in intron RNA abundance between glucose- and ethanol-grown cultures, this data therefore suggests condition-dependent group II intron processing or stabilization by RNA-processing enzymes, which might respond to glucose repression. The underlying molecular mechanism remains to be explored.

While accumulation of introns in glucose-grown cultures might be a mere side effect of condition-dependent intron turn-over rates, an interesting fundamental question is whether these group II introns have an impact on mitochondrial functions. In yeast, it has been demonstrated that efficient splicing of group I introns plays a regulatory role and that accumulation of group I introns can result in ‘exon-reopening’, hindering mitochondrial function (64,96,108), while in higher eukaryotes, non-coding mtRNA may play a role in cell signalling and mitochondrial expression (93,94). Such functions have not yet been described for group II introns, and the impact of lariats on yeast physiology is controversial.

The group II introns are not conserved within the *Saccharomyces* genus (109), and as intron-free strains did not show a different phenotype in ethanol medium, it was assumed that group II introns are phenotypically neutral (85,86). Conversely, one study reported increased respiration rates and mitochondrial volume during growth on glucose medium using a fully intron-less *S. cerevisiae* strain, a phenotype which could also be achieved by overexpressing the Mss116p RNA-helicase and –chaperone involved in group II introns splicing (31,96).

Intron-less strains from the CEN.PK *S. cerevisiae* lineage (28), used for transcriptome analysis in the present study, are unfortunately not available and are challenging to construct. The response of an intron-less strain to growth on glucose and ethanol was therefore performed with an available strain from a different lineage. Unfortunately, although small changes in physiology were observed, results presented in earlier studies could not be reproduced (96), which might be attributed to different growth conditions (*e*.*g*. medium composition). Nevertheless, no substantial physiological difference could be observed between intron-less and control strain grown with either glucose or ethanol as carbon source (Figure 6). There is no transcriptome data available for the 161 lineage control during growth on glucose and ethanol, there is therefore no evidence of condition-dependent variation in group II introns levels in this specific lineage. More research will therefore be required by either constructing intron-less CEN.PK strains or by quantifying mtRNA levels in strains of the 161 lineage, to conclude on a potential physiological effect of group II introns.

Little is known about the context-dependency of mitochondrial gene expression and splicing. Using the developed method, the impact of environment on transcriptional responses in the tractable model organism *S. cerevisiae*, can be uncovered at a base pair-level resolution, both in terms of RNA quantity and processing under different conditions. For instance, mitochondrial activity can be easily tuned between ‘minimal service’ in anaerobic cultures with rich medium, to ‘hyperactivity’ in respiratory, unrepressed growth conditions (e.g. the ethanol-fed aerobic cultures used in the present study). Considering the important role of mitochondria in aging in mammals, monitoring transcriptomic responses of yeast mitochondria to oxidative stress or in chronologically aging cultures might reveal new layers in gene expression regulation. The presented methodology therefore opens the door to a deeper understanding of mitochondrial processes and their regulation.

## Supporting information

SI_Figures-Tables-MM-sequences

SI_Readcounts

## Data availability

R scripts and CEN.PK113-7D mitochondrial reference sequences are deposited and can be accessed freely at https://gitlab.tudelft.nl/charlottekoste/mitornaseq. All raw sequencing data is deposited at NCBI (www.ncbi.nlm.nih.gov): RNA-seq data and normalized expression levels are deposited at GEO under accession number GSE219013 and DNA sequencing data as well as *de novo* assemblies are deposited under BioProject ID PRJNA902953.

## Supplementary data

SI_Figures-Tables-MM-sequences.pdf – Contains supplementary figures, supplementary tables, additional Materials and methods and raw RNA and DNA sequences used in the text.

SI_readcounts.xlsx – Contains a spreadsheet with raw read counts per gene, gene counts normalized to CPM and gene counts normalized to CPM and mtDW.

## Acknowledgements

The authors thank Prof. Alan Lambowitz and dr. Georg Mohr for providing strains 161-U7 and 161-I^0^ and Prof. Alan Lambowitz for feedback on the manuscript.

## Author contributions

CCK, AK, PDL and JMD conceptualized the study and provided scientific input, CCK, AK and ML designed and performed experiments, AK and MvdB designed the bioinformatics pipeline, CCK, AK and MvdB performed data analysis and CCK, AK and PDL wrote the manuscript. All authors read and agreed to the final manuscript.

## Footnotes

1 https://international.neb.com/faqs/2011/11/28/is-the-em-e-coli-em-poly-a-polymerase-able-to-elongate-short-and-surface-immobilized-oligoribonuc

