## Supplementary material for "Long-read direct RNA sequencing of the mitochondrial transcriptome of *Saccharomyces cerevisiae* reveals condition-dependent intron turnover": SI_Figures-Tables-MM-sequences

Koster *et al.*

This supplementary Information contains:

### Supplementary Figures

#### Figure S1

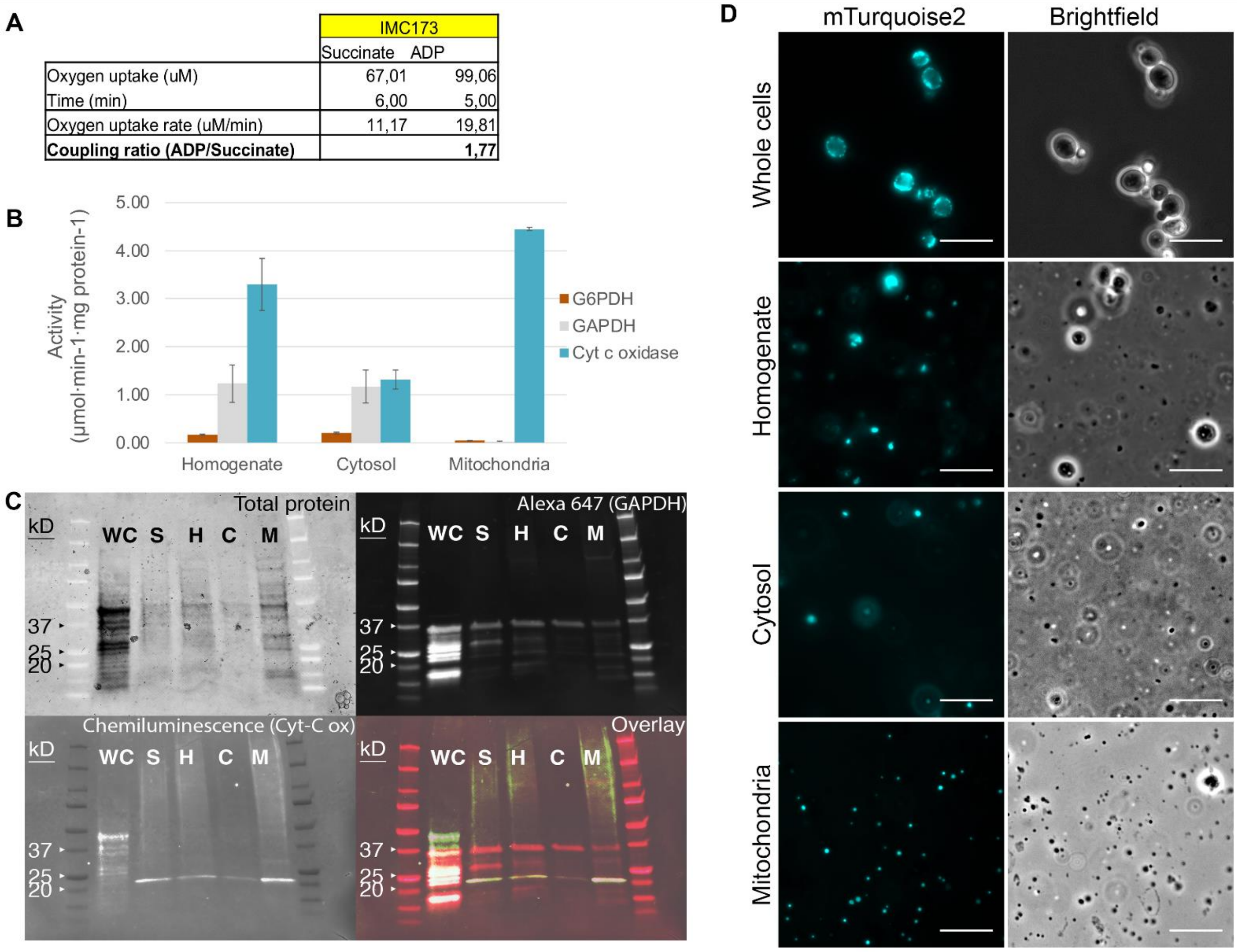

Figure S 1. **Quality assessment of the mitochondria isolation protocol using strain IMC173 (preSU9-mTq2, blue fluorescent mitochondria)**. A) Oxygen consumption rate of isolated mitochondria supplied first with succinate then with ADP. Details on the experimental setup are described in the supplementary methods. The ratio of the oxygen uptake rates before and after the ADP addition informs on the activity of the respiratory chain and therefore on the integrity of the mitochondrial membranes. The coupling (P/O) ratio when using succinate as substrate is expected to be close to 1.5 (1). B) Enzyme activity of the cytosolic glucose-6-phosphate dehydrogenase (G6PDH) and glyceraldehyde-3-phosphate dehydrogenase (GAPDH) and mitochondrial Cytochrome *c* oxidase (Cyt *c*, mitochondrial). Activity was measured in different fractions obtained throughout the mitochondria isolation protocol, homogenate: lysed spheroplasts, cytosol: cytosolic fraction after differential centrifugation, mitochondria: mitochondrial fraction after differential centrifugation. C) Western Blots of the mitochondrial fraction. GAPDH (cytosolic, 35,7 kDa) was detected with conjugated anti-GAPDH antibodies containing Alexa 647 and Cytochrome *c* oxidase subunit 3 (mitochondrial, 30 kDa) was detected with an anti-Cox3 antibody and a secondary antibody with an HRP group and chemiluminescent substrate. Protein content was imaged using BioRad Stain-Free gels activated under UV. Different fractions of the mitochondrial isolation protocol were analyzed, WC: whole cells, S: supernatant after spheroplasting, H: homogenate (lysed spheroplasts), C: cytosolic fraction, M: Mitochondrial fraction. D) Microscopy images of whole cells, homogenate, cytosol and mitochondria, left: blue fluorescence (ex: 436 nm, em: 480 nm), right: widefield image.. Scale bars represent 10 µm.

#### Figure S2

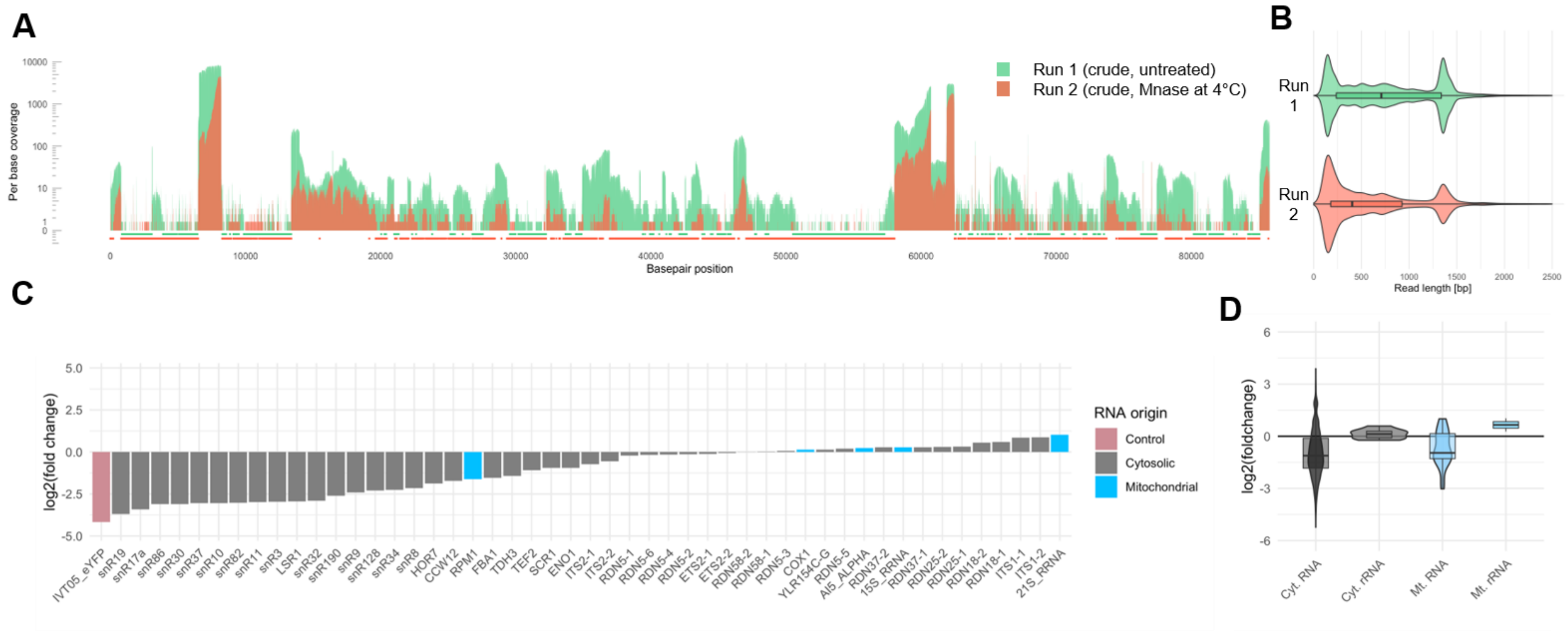

**Figure S 2. Comparison of sequencing data of crude mitochondrial fraction (run 1) and MNase-treated mitochondria (run 2)**. A) Per-base-coverage of the full mitochondrial genome, with per-base coverage of crude mitochondria (run 1) in green and per-base coverage of MNase-treated mitochondria (run 2) in orange on $\log_{10}$ scale. B) Distribution of obtained read lengths for run 1 and run 2 shown as violin plots, the mean is shown as a box plot where the outer ends of the box represent the 25$^{th}$ and 75$^{th}$ percentile of the data. C) Top 50 most abundant transcripts of run 2 and their fold change of CPM between run 1 and run2, sorted by $\log_2$ fold change in increasing manner. D) $\text{Log}_2$ fold change of CPM of run 1 and 2, grouped by different type (cytosolic RNA except rRNA, cytosolic rRNA, mitochondrial RNA except rRNA and mitochondrial RNA).

#### Figure S3

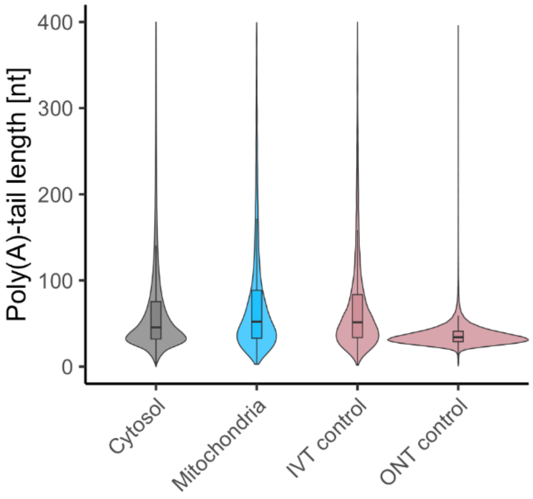

**Figure S 3. Estimated Poly(A)-tail lengths after enzymatic polyadenylation.** The length of the poly(A)-tail was estimated by Nanopolish-PolyA on the obtained RNAseq data for RNA of mitochondrial (no native poly(A) tail) and cytosolic origin (natively poly(A)-tailed) and of in vitro-synthesized control RNA (no native poly(A)-tail). The length of the poly(A)-tail of the ONT control RNA (not enzymatically polyadenylated, contains poly(A)-tail) was determined as a control. Distribution of poly(A)-tail lengths is shown as violin plots, the mean is shown as a box plot where the outer ends of the box represent the 25$^{th}$ and 75$^{th}$ percentile of the data.

#### Figure S4

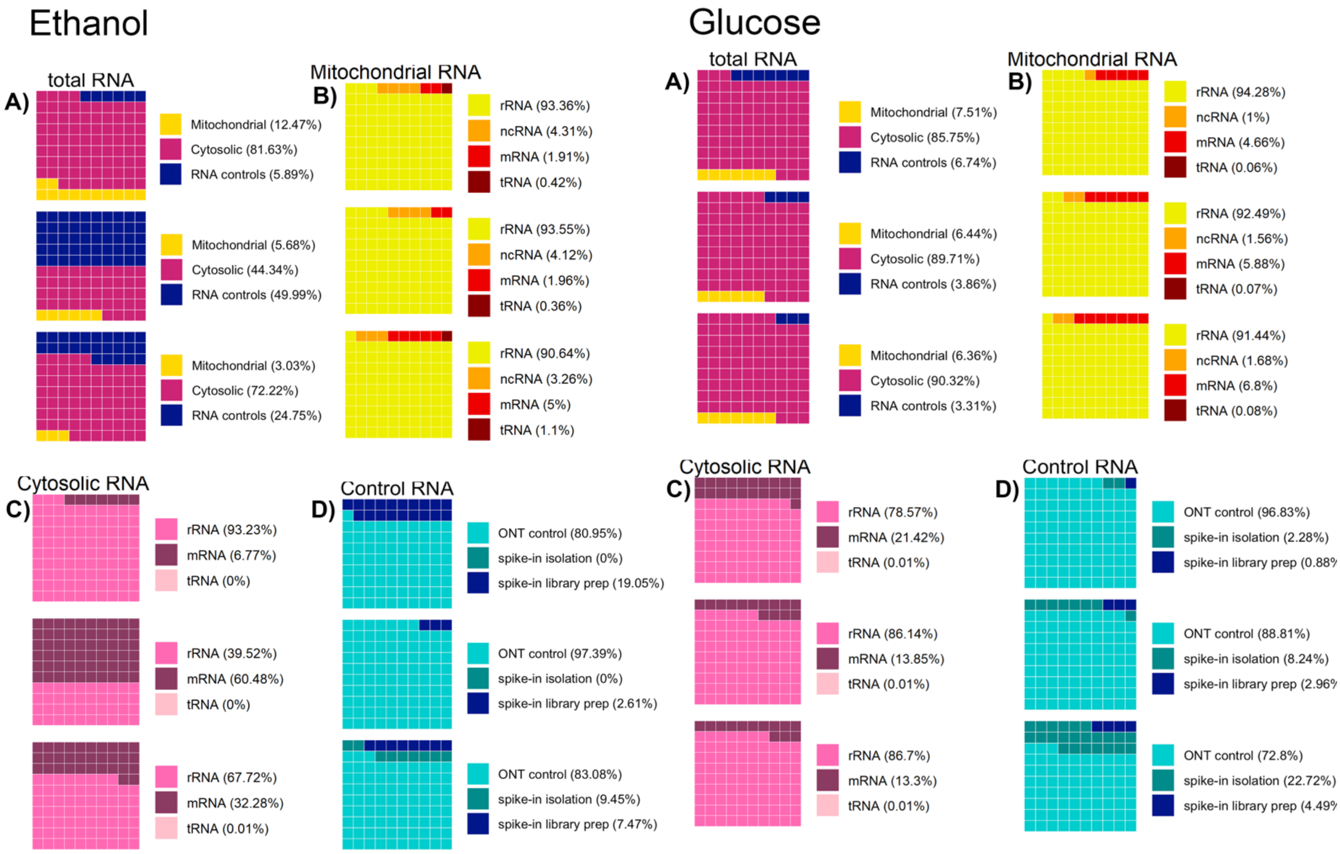

**Figure S 4. Waffle plots of read origin and type for the different sequencing runs of triplicate experiments on glucose and ethanol**. A) Distribution of the spatial origin of sequencing reads between mitochondrial, cytosolic and control RNA. Breakdown of the distribution of different types of mitochondrial RNA (B), cytosolic RNA (C), and spiked-in control RNA (D). One square represents 1/100 of the total number of reads. Left panel: glucose-grown cultures, right panel: ethanol-grown cultures

#### Figure S5

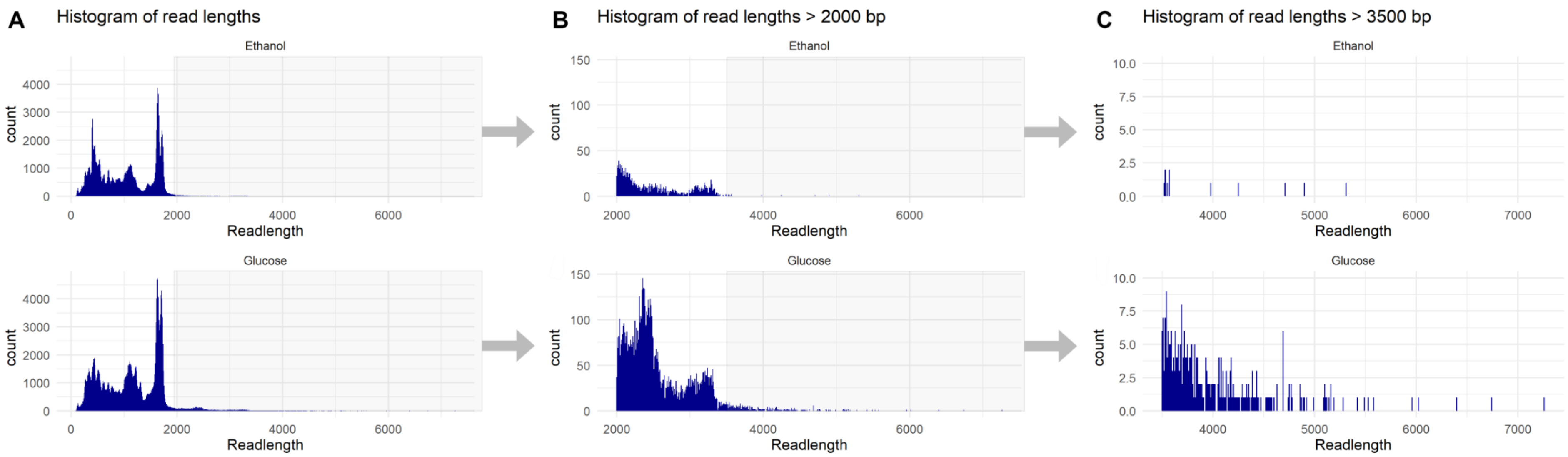

**Figure S 5. Distribution of read counts according to read length in the sequencing dataset**. The data show the cumulated read lengths obtained in three replicate experiments on either glucose (top graphs) or ethanol (bottom graphs) as carbon source. A) Distribution of all read lengths. B) and C): Zoom in for the distribution for read lengths larger than 2000 bp (B) and 3500 bp (C) that have lower abundance.

#### Figure S6

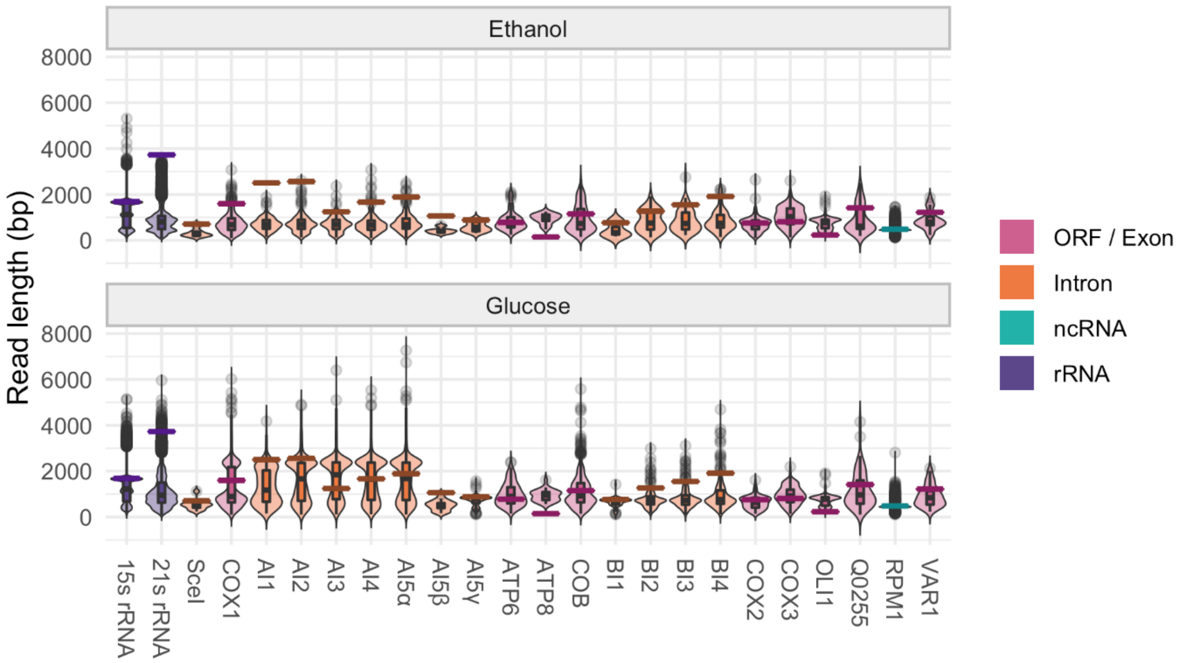

**Figure S 6. Read length distribution per gene**. The distribution of the average read length of the r=triplicate experiments is shown as violins, the average mean read length is shown as a box plot where the outer ends of the box represent the 25th and 75th percentile of the data. Outliers are shown as circles. The gene length is indicated with a horizontal line. Reads are colored per type, mRNA (pink), ncRNA (green) rRNA (purple). Spliced genes are separated between the main exon (pink) and the different spliced-out introns (orange).

Figure S7

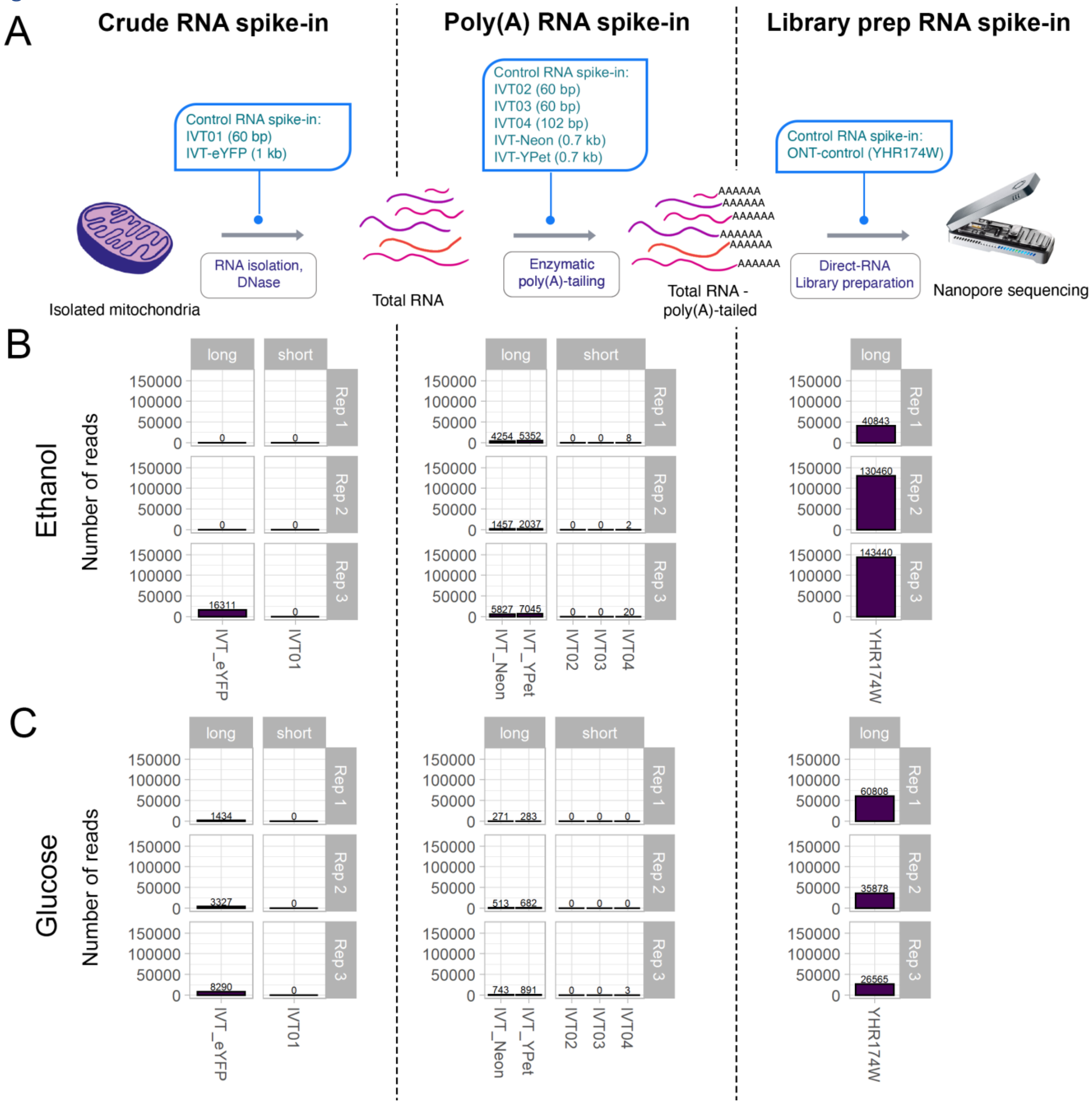

**Figure S 7. Overview of control RNA abudance over all transcriptome experiments**. **Number of control reads sequenced for all sequenced replicates.** A) Schematic overview of the mtRNA sequencing protocol indicating in which steps the control RNAs are spiked. All reads were spiked in equimolar amounts (45 fmol), except for the ONT control read YHR174W which was spiked at a concentration of 61 fmol according to the ONT direct sequencing protocol. B) and C) Number of sequenced reads for the controls spiked as indicated in A). Data are shown for each independent culture replicate (Rep 1 to Rep 3) and for each spiking step..The absolute number of passed reads in each replicate is plotted without normalization, the numbers above the bars show the number of passed reads per control sequence. 'Long' denotes a read > 150 bp, 'short' reads < 150 bp. Rep = Replicate. B) Cultures grown on ethanol, C) cultures grown on glucose.

#### Figure S8

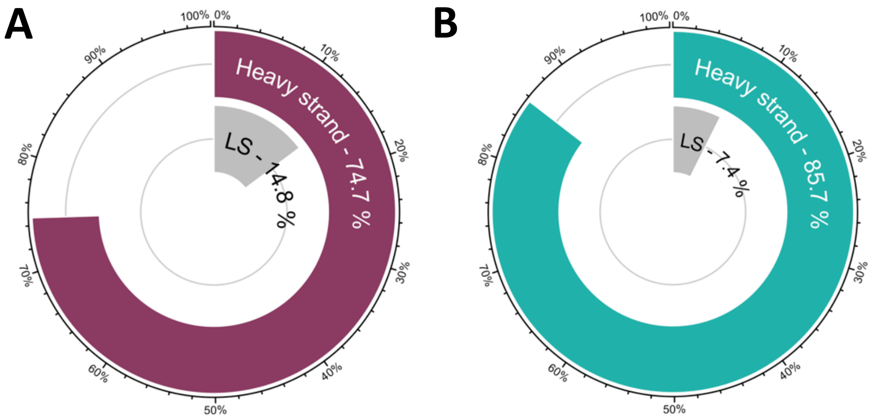

**Figure S 8. Breadth of coverage of the heavy and light strand of the mitochondrial genome**. A) Ethanol-grown cultures . B) glucose-grown cultures. The coverage is defined as the number of bases that were significantly expressed in triplicate experiments, divided by the size of the mitochondrial genome.

Figure S9

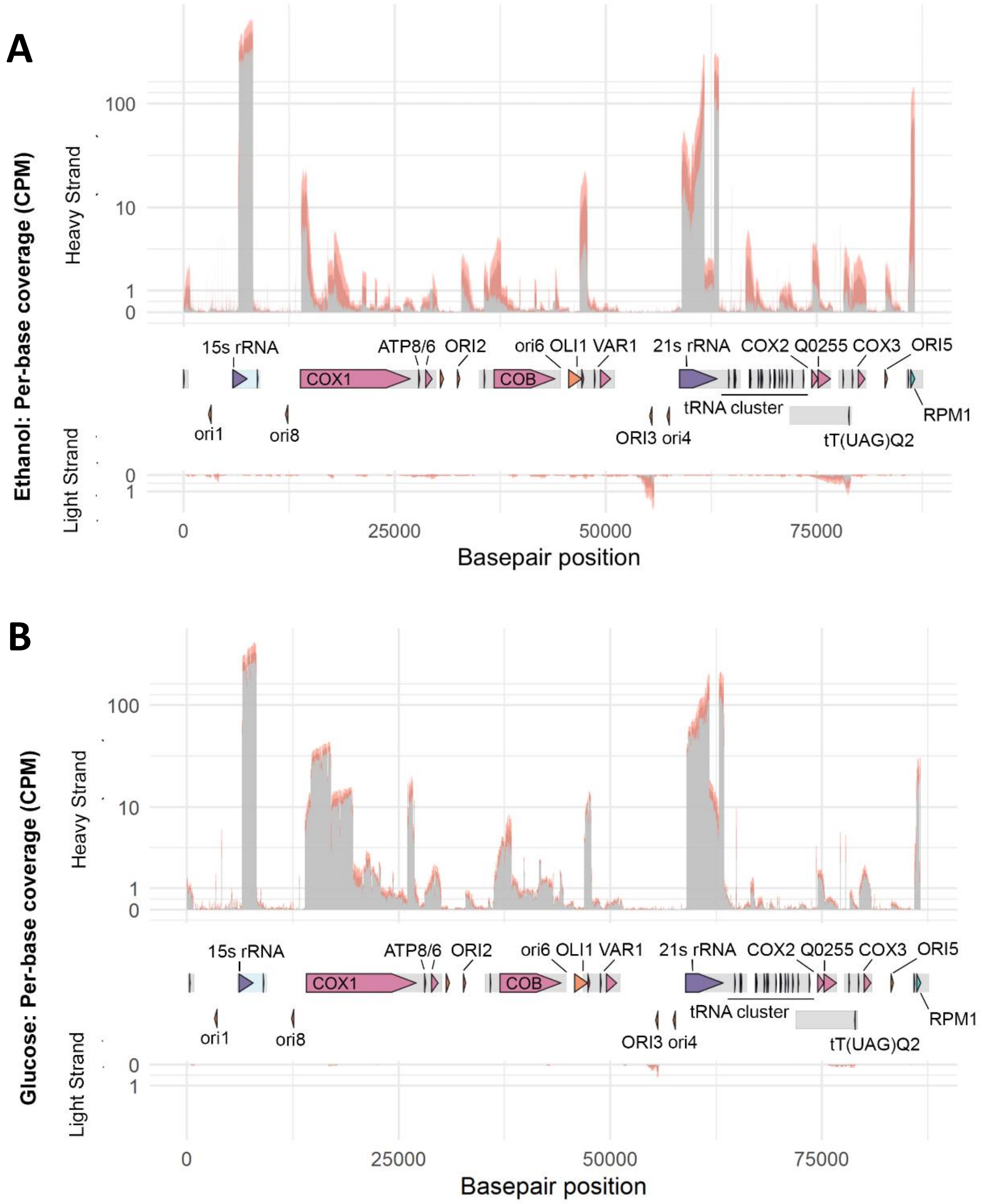

Figure S 9. Per-base coverage of the mitochondrial transcriptome. A) Ethanol-grown cultures B) glucose-grown cultures. bottom). The data are normalized per million bases as Counts-Per-Million (CPM) and both the heavy (forward, top graph) and light (reverse, bottom graph) strands of the mitochondrial DNA are represented. Coverage depth is represented in grey, while the standard deviation between the sequencing depth of triplicate experiments is shown as a red area. The mitochondrial genome is shown between the graphs. Grey boxes indicate primary transcripts.

Figure S10

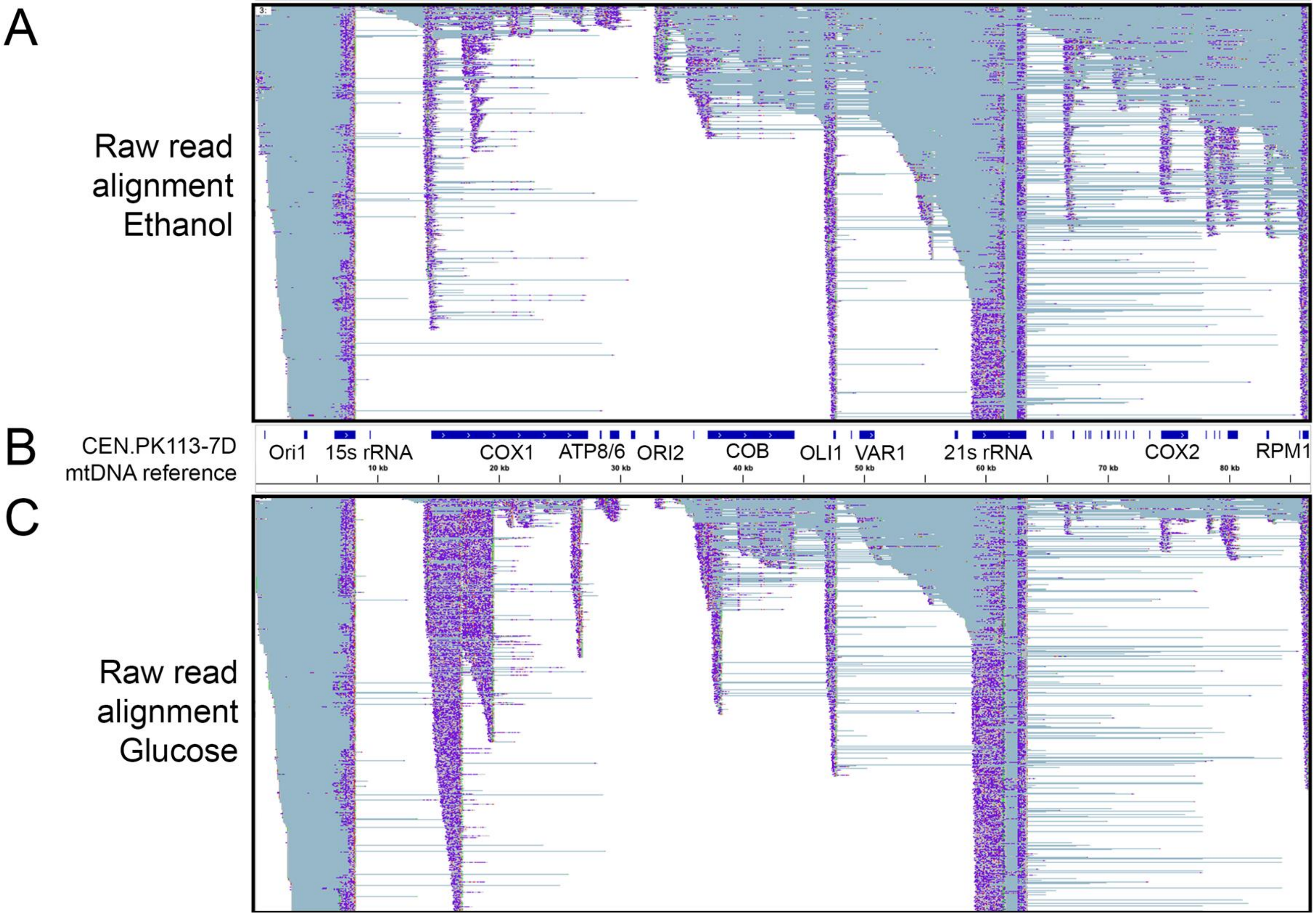

**Figure S 10. Raw read alignment in igv of mitochondrial RNAseq data across the entire mitochondrial genome.** A single representative replicate is shown for growth on ethanol (A) and growth on glucose (C). **Each line is one read, colors indicate the different bases of the read (A = green, T = red, C = dark blue, G = orange). Grey-blue lines indicate gaps in the reads.** B) Annotation of the CEN.PK113-7D mitochondrial reference genome corresponding to the read alignments shown in A) and C).

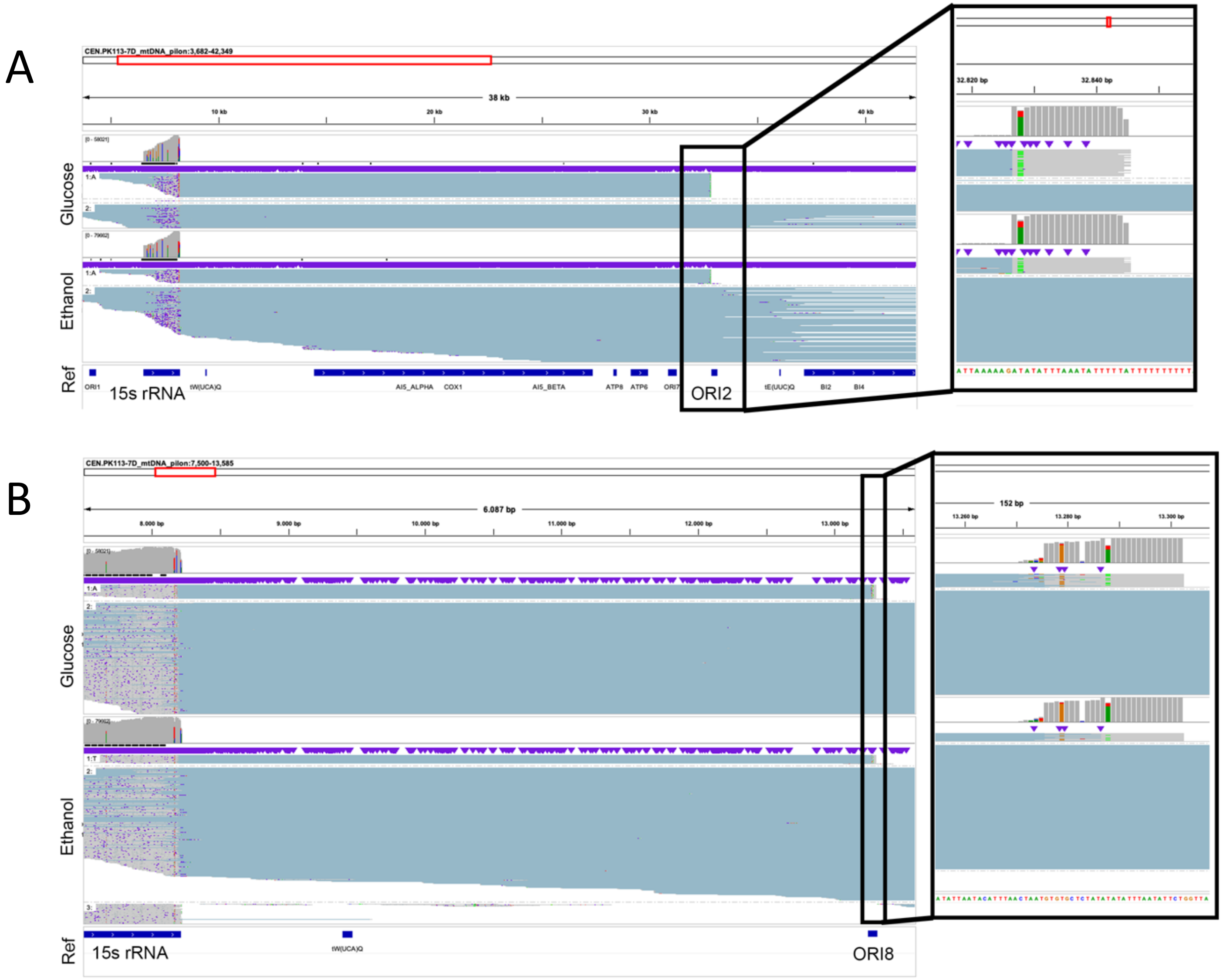

**Figure S 11. Raw read alignment in igv of mitochondrial RNAseq data in the 15s rRNA to *ORI2* (A) and *ORI8* (B) loci**. In each panel a single representative replicate is shown for ethanol (top graph) and glucose-grown culture (bottom graph). Grey, dark blue, yellow, green and red bases indicate read mapping, a light blue line indicates a gap in the read. The annotation of the CEN.PK113-7D mitochondrial reference genome is shown at the bottom of each panel.

Figure S12

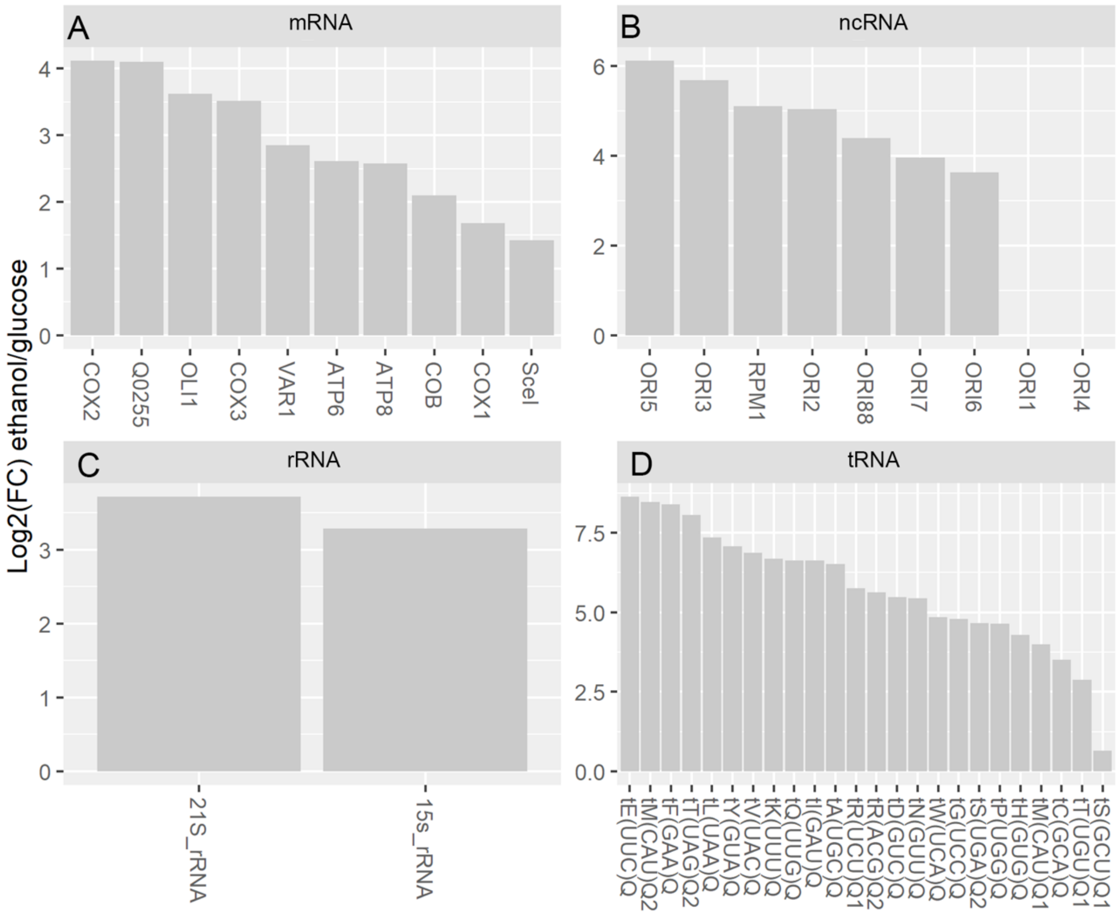

Figure S 12. **Change in expression of mitochondrial RNA species between glucose- and ethanol-grown cultures.** Data represent the *log2* fold-change (FC) of transcript expression of mitochondrial RNA in ethanol as compared to glucose cultures. A) mRNA (no separation made between introns and exon reads), B) ncRNA, C) rRNA, D) tRNA

Figure S13

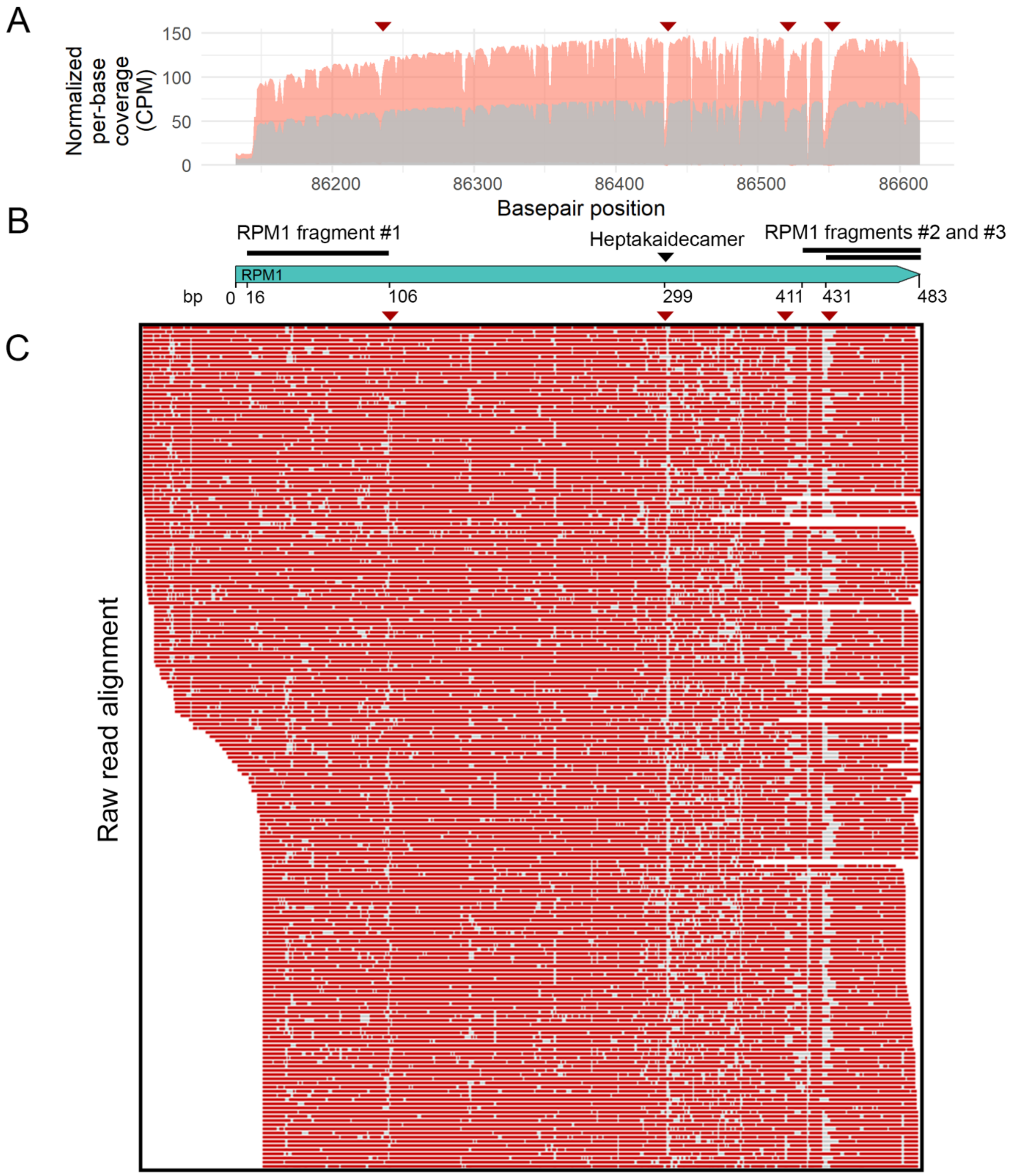

**Figure S 13. Coverage plot and read alignment at the *RPM1* locus.** B) Schematic representation of the *RPM1* locus, including the RNA fragmentation pattern (black boxes) and heptakaidecamer location (black arrow) proposed by Turk *et al.* (2). A) Average coverage of triplicate experiments of the gene normalized to CPM on ethanol (grey). The standard deviation of coverage is shown in red. Decreases in coverage caused by RNA processing are indicated with red arrows. C) Visualization of the sequencing reads mapped to the *RPM1* locus using Tablet. Red lines indicate correct mapping of sequencing reads to the location on the mtDNA, grey lines indicate a gap in the reads. Gaps potentially resulting from RNA processing are indicated with red arrows.

Figure S14

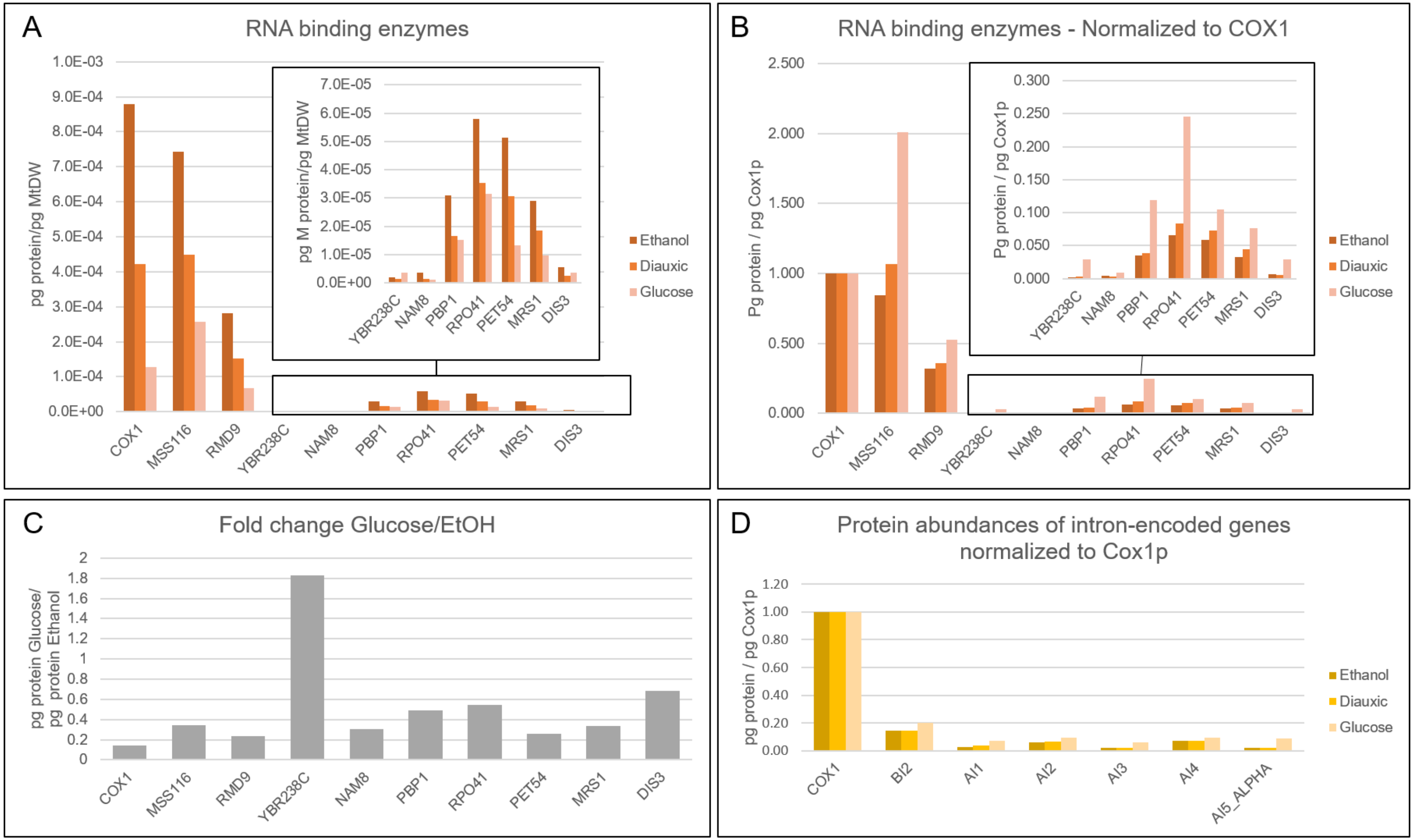

**Figure S 14. Abundance of (putative) mitochondrial RNA binding proteins as determined by (3), and the abundance of intron-encoded proteins**. A-C) From the datasets from Mitchell *et al* (4) Tsvetanova *et al* (5) of RNA-binding proteins in yeast, a list of RNA-binding proteins was extracted. This list was cross-referenced with the mitochondrial proteome a published by Di Bartolomeo *et al.* to identify a list of nine (putative) mitochondrial

RNA binding proteins. All panels in this figure represent protein abundance as reported by di Bartolomeo and colleagues (6). A) Absolute level of the putative RNA-binding proteins and Cox1p in pg mitochondrial protein normalized by mitochondrial dry weight during growth on ethanol and on glucose and during the diauxic shift. B) Relative abundance of the nine putative RNA binding proteins normalized to the level of Cox1p. C) Fold-change in RNA binding proteins abundance between cultures with glucose and ethanol as carbon source. D) Abundance normalized to Cox1p of intron-encoded proteins during growth on ethanol and on glucose and during the diauxic shift.

Figure S15

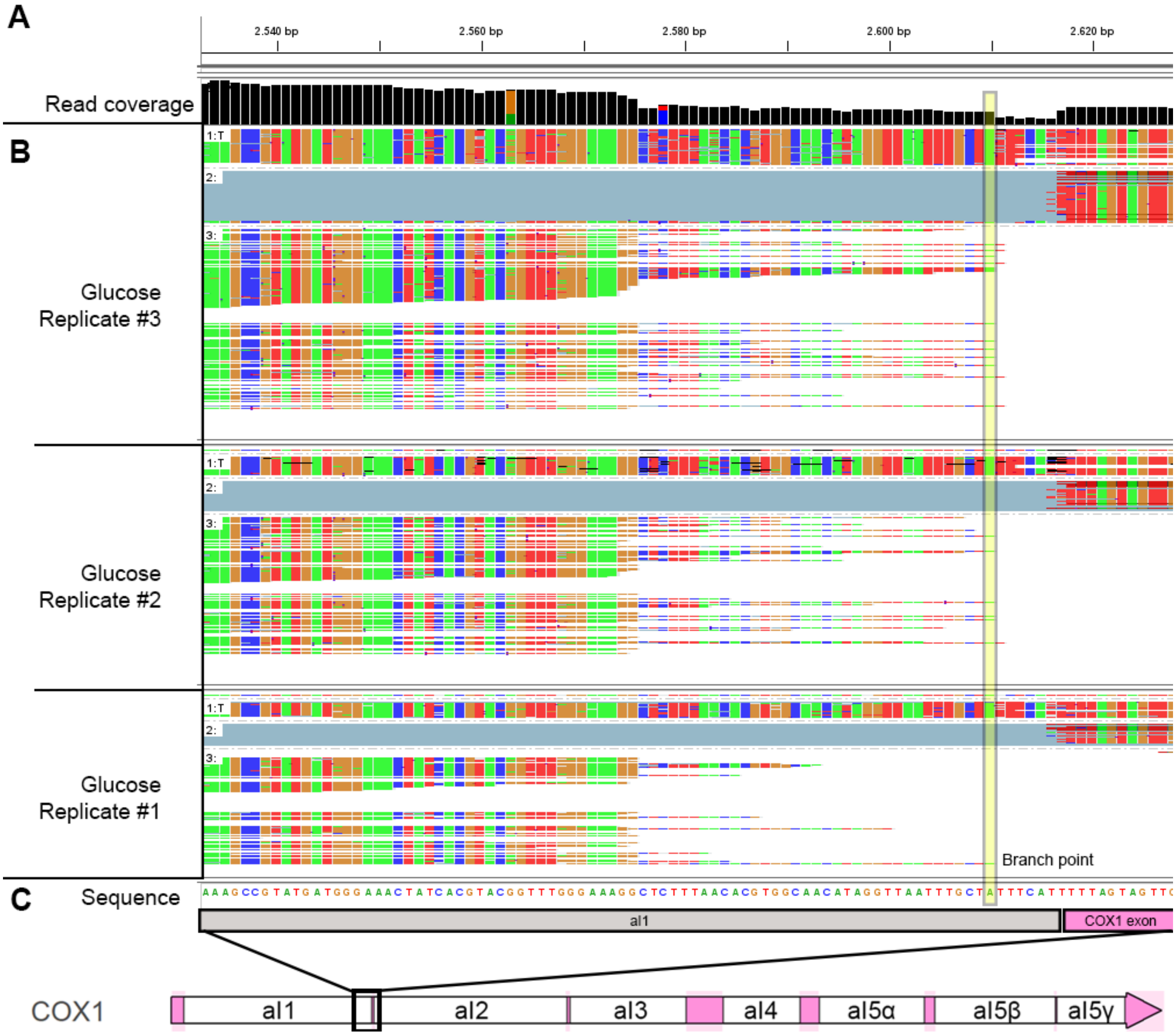

**Figure S 15. Read mapping at the 3’ branch point of *COX1* aI1 intron sequences**. A) Coverage plot (in black) of the locus. B) Mapping of raw reads for three separate replicates mapped to the reference sequence. Each line is one read, colors indicate the different bases of the read (A = green, T = red, C = dark blue, G = orange). Grey-blue lines indicate gaps in the reads. In A) and B) the branch point as described by Yang and colleagues (where the circularization of the RNA through a covalent bond occurs, (7)) is indicated with a yellow box. C) Schematic overview of the DNA sequence of the COX1-aI1 intron-exon junction.

Figure S16

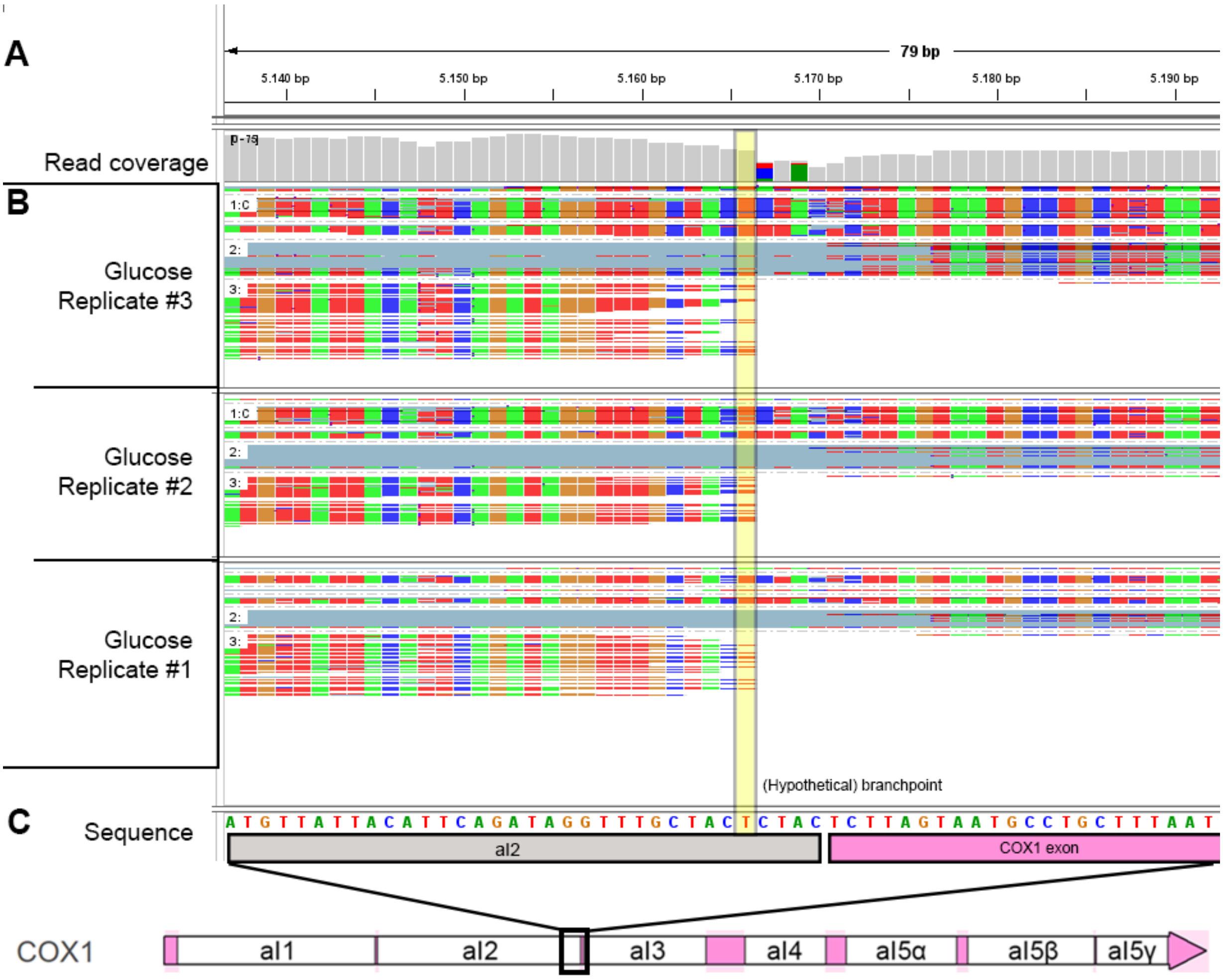

**Figure S 16. Read mapping at the 3' branch point of *COX1* aI2 intron sequence.** A) Coverage plot (in grey) of the locus. B) Mapping of raw reads for three separate replicates mapped to the reference sequence. Each line is one read, colors indicate the different bases of the read (A = green, T = red, C = dark blue, G = orange). Grey-blue lines indicate gaps in the reads. The exact branch point of aI2 is not described in literature, therefore the branchpoint was hypothesized to be the Uracil-residue within a palindromic sequence where reads end, as indicated with a yellow box in A) and B). C) Schematic overview of the DNA sequence of the COX1-aI2 intron-exon junction.

Figure S17

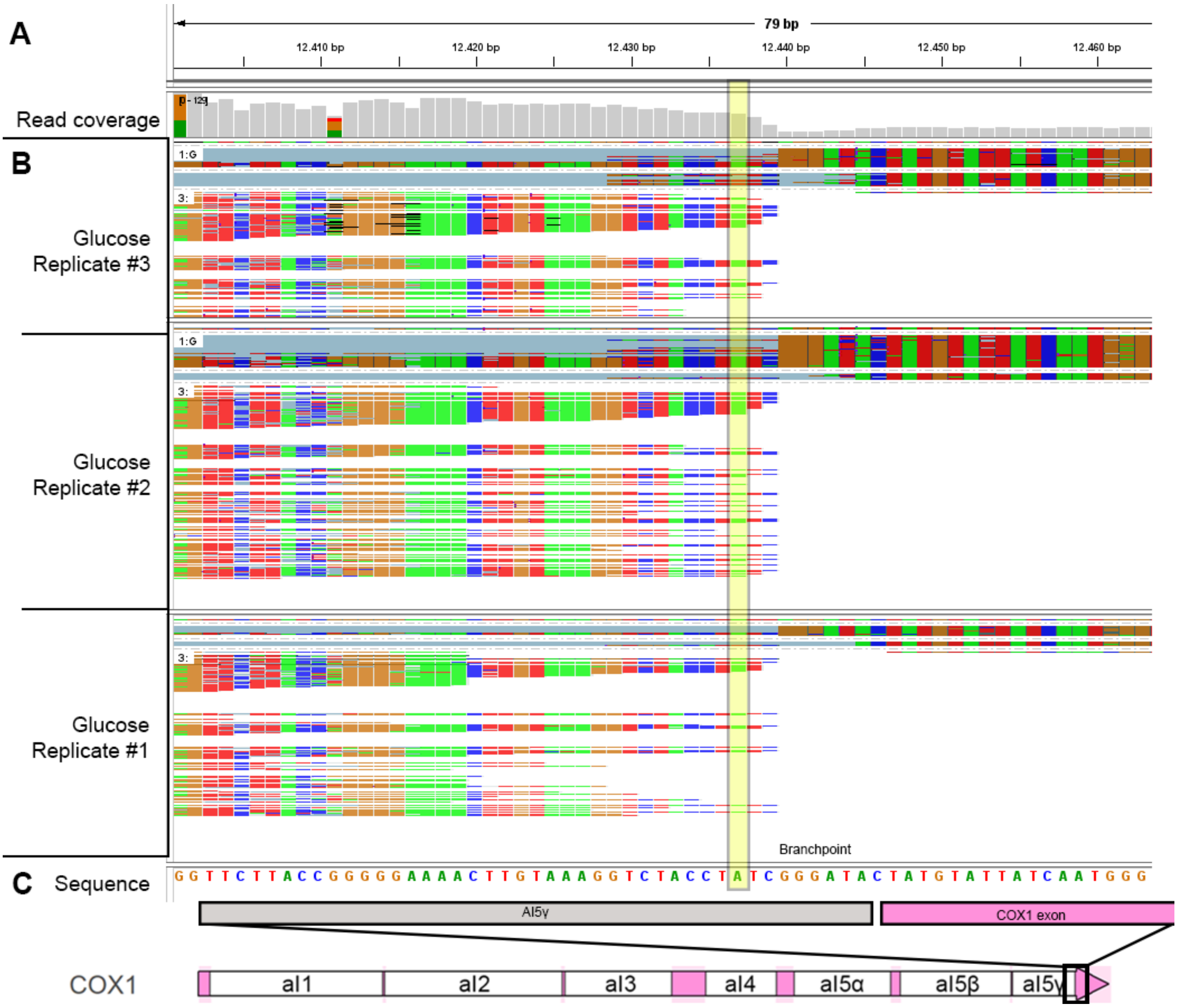

**Figure S 17. Read mapping at the 3' branch point of *COX1* aI5γ intron sequence**. A) Coverage plot (in gray) of the locus. B) Mapping of raw reads for three separate replicates mapped to the reference sequence. Each line is one read, colors indicate the different bases of the read (A = green, T = red, C = dark blue, G = orange). Grey-blue lines indicate gaps in the reads. In A) and B) the branch point as described by Schmelzer and Schweyen (where the circularization of the RNA through a covalent bond occurs, (8)) is indicated with a yellow box . C) Schematic overview of the DNA sequence of the COX1-aI5γ intron-exon junction.

Figure S18

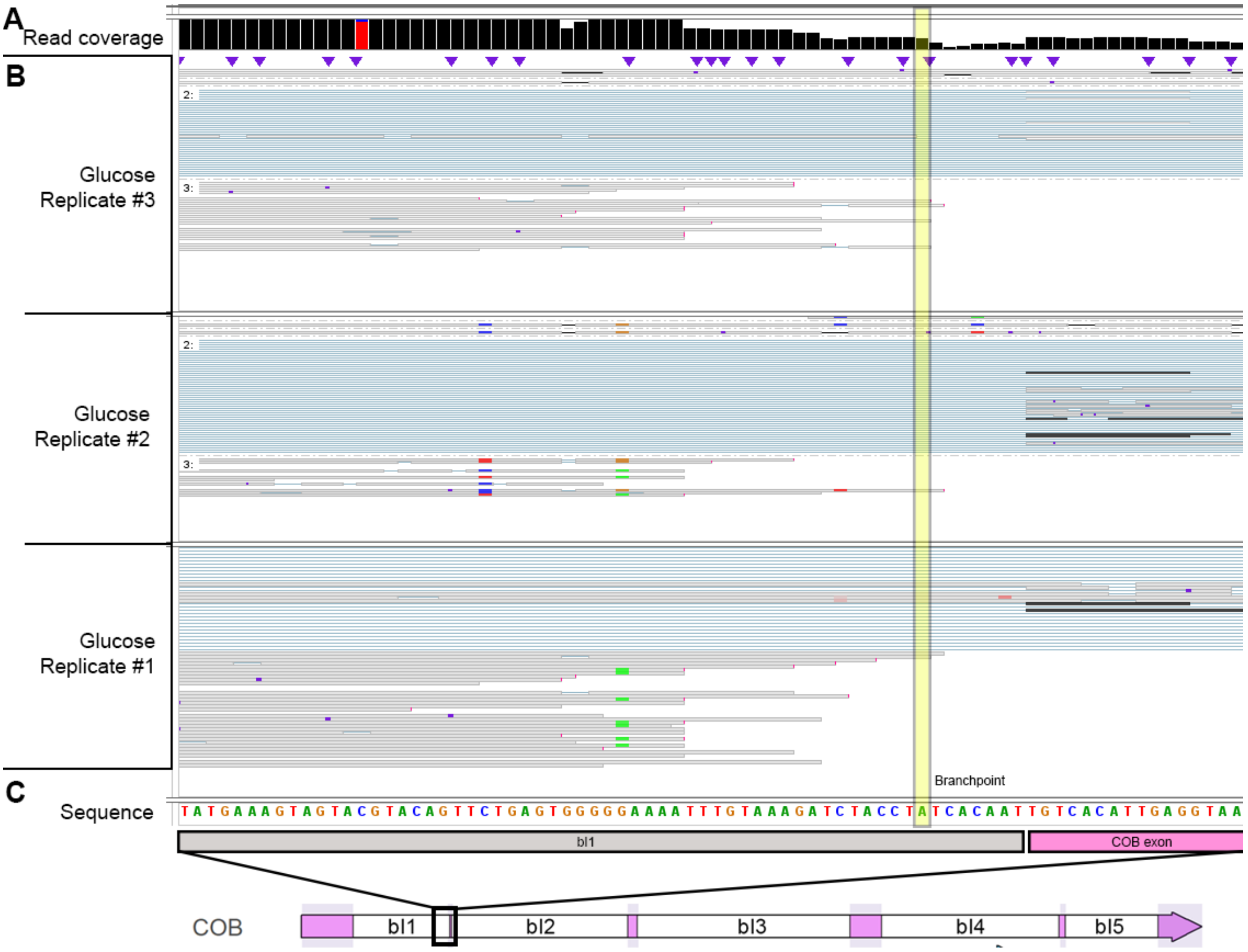

**Figure S 18. Read mapping at the 3’ branch point of *COX1* bI1 intron sequence.** A) Coverage plot (in black) of the locus. B) Mapping of raw reads for three separate replicates mapped to the reference sequence. Each grey line is one read, blue lines indicate gaps in the reads. In A) and B) the branch point as described by Schmelzer and Schweyen (where the circularization of the RNA through a covalent bond occurs, (8)) is indicated with a yellow box. C) Schematic overview of the DNA sequence of the COX1-bI1 intron-exon junction.

#### Figure S19

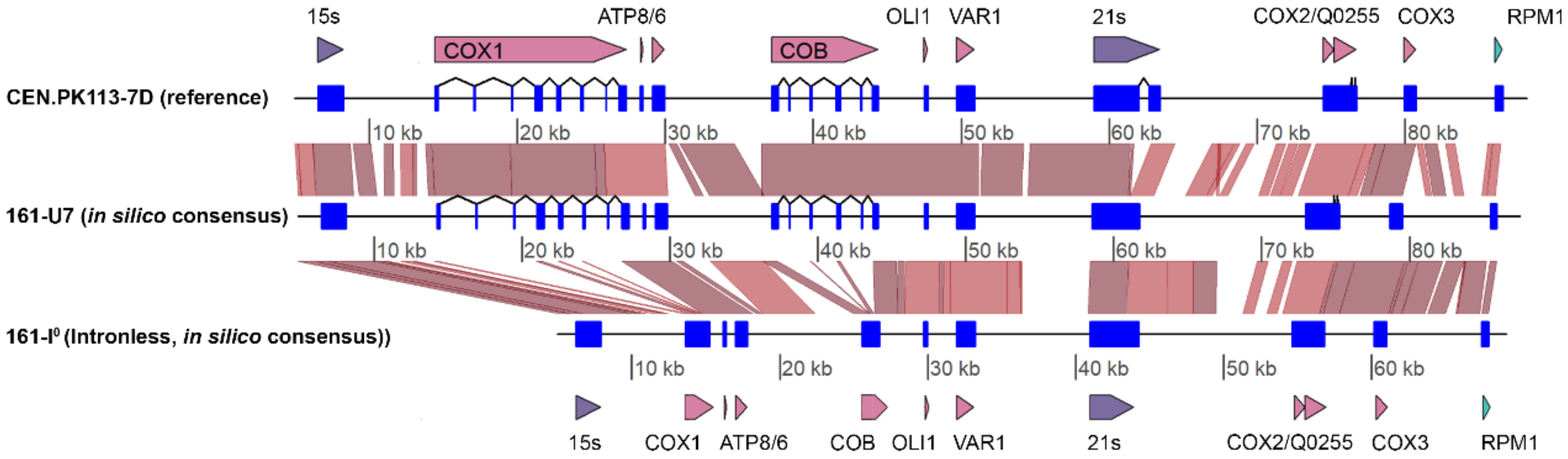

**Figure S 19. Alignment of the mitochondrial genome of CEN.PK113-7D, and consensus sequences of the mitochondrial genomes of 161-U7 and 161-I$^0$.** The mitochondrial genome is represented as a black line and the respective coordinates are indicated below each genome/contig. The locations of exon mRNA, rRNA and ncRNA are indicated as blue squares, tRNA and ori are not shown. Spliced exons are connected by a line, annotations of the sequences are indicated above (CEN.PK113-7D, 161-U7) and below (161-I$^0$) the alignment. Red boxes or lines indicate a sequence identity of > 95 % (BLAST) between the two alignments.

### Supplementary Tables

#### Table S1

**Table S 1. Additional strains used in this study**

| Strain name | Relevant genotype | Parental strain | Reference |
|---|---|---|---|
| IMX1714 | *MATa sga1*::*Spycas9*-natNT2 | CEN.PK113-7D | (9) |
| IMX2100 | *MATa sga1*::*Spycas9*-natNT2 *Δura3* | IMX1714 | This study |
| IMX2219 | *MATa sga1*::*Spycas9*-natNT2 *Δura3 tom70::TOM70*-6xHis | IMX2100 | This study |
| IMC173 | *MATa sga1*::*Spycas9*-natNT2 *Δura3 tom70::TOM70*-6xHis pUDC329 (*preCOX4-mTq2, URA3*) | IMX2219 | This study |

#### Table S2

Table S 2. Plasmids used in this study

| Plasmid expression and construction | | | |
|---|---|---|---|
| Plasmid name | Relevant plasmid genotype | Purpose of plasmid | Source |
| pUDC329 | *pTEF1-preSU9-ymTq2-tENO2 URA3 CEN6/ARS4 bla, ori* | Expression mitochondrial mTq2 | This study |
| pUD538 | *URA3 CEN6/ARS4 bla, ori ColE1* | GG drop-out backbone | (10) |
| pUD585 | *pTEF1 YTK-part 2, CamRi, ColE1* | GG part 2 | (11) |
| pGGkp320 | *preSU9 YTK-part3a, CamR, ColE1* | GG part 3a | This study |
| pGGkp308 | *ymTurquoise2 YTK-part3b, CamR, ColE1* | GG part3b | This study |
| pYTK055 | *tENO2 YTK-part4b, CamR, ColE1* | GG part 4 | (12) |
| pDRF1-GW ymTurquoise2 | *ymTurquoise2, URA3, 2μ, bla, ColE1* | Template for pGGkp308 | (13) |
| pFA6a-link-ymNeongreen-URA3 | *ymNeongreen URA3 2μ, bla, ColE1* | Template for pGGkp304 | (13) |
| pDRF1-GW ymYPET | *ymYPET URA3 2μ, bla, ColE1* | Template for pGGkp305 | (13) |
| **Cas9 editing** | | | |
| Plasmid name | Relevant plasmid genotype | Purpose of plasmid | Source |
| pUDR107 | *URA3-gRNA* hphNT1 *2μ, bla, ColE1* | Cas9-cleavage of *URA3* | (14) |
| pUDR621 | *TOM70-gRNA URA3, 2μ, bla, ColE1* | Cas9-cleavage of *TOM70* | This study |
| pUD999 | *TOM70-6xHis repair fragment, bla, ColE1* | Repair fragment template | This study |
| **Templates used for in vitro RNA synthesis** | | | |
| Plasmid name | Relevant plasmid genotype | Purpose of plasmid | Source |
| pGGkp304 | *ymNeonGreen YTK-part3, CamR, ColE1* | Template for In vitro RNA synthesis | This study |
| pGGkp305 | *ymYPET YTK-part3, CamR, ColE1* | Template for In vitro RNA synthesis | This study |
| **meYFPco-LL-spinach** | *YFP, bla, ori, ColE1* | Template for In vitro RNA synthesis | (15) |

### Table S3

#### Table S 3. Primers used in this study

| # ID | PURPOSE | SEQUENCE[1] |
|---|---|---|
| **10900** | Colony PCR / | GTTCTTTCCTGCGTTATCCC |
| **10901** | Sanger sequencing pUDC329 | AAGTCTGTGCTCCTTCCTTC |
| **17422** | Sanger sequencing | TTCTCTGTAAGCGGTGAAGG |
| **17423** | pUDC329 | TGATCCCAGCAGCAGTAACG |
| **17713** | Attaching GG-flanks | GCAT**CGTCTC**ATC**GGTCTC**ATATGATGGCTTCTACTAGAGTTTTGG |
| **17714** | to *preSU9* | ATGC**CGTCT**CA**GGTCTC**AAGAACCAGCTCTCTTTTGGAAAGCTT |
| **17381** | Attaching GG-flanks | GCAT**CGTCTC**ATC**GGTCTC**ATTCTGTTAGTAAAGGTGAAGAATTGTT |
| **17706** | to *ymTurquoise2* | ATGC**CGTCTC**A**GGTCTC**AGGATCCTTTATACAATTCATCCATACC |
| **17383** | Attaching GG-flanks | GCAT**CGTCTC**ATC**GGTCTC**ATATGGTCTCTAAGGGTGAAGAAGATAACATGG |
| **17707** | to *ymNeonGreen* | ATGC**CGTCTC**A**GGTCTC**AGGATCCCTTGTACAATTCGTCCATACCC |
| **17386** | Attaching GG-flanks | GCAT**CGTCTC**ATC**GGTCTC**ATATGGTCTCCAAGGGCGAAGAATTGTTCACC |
| **17708** | to *ymYPET* | ATGC**CGTCTC**A**GGTCTC**AGGATCCTTTATACAATTCATCCATACC |
| **15155** | TOM70 gRNA | TGCGCATGTTTCGGCGTTCGAAACTTCTCCGCAGTGAAAGATAAATGATCAATTCTTTGTTGAACTTTAGGTTTTAGAGCTAGAAATAGCA AGTTAAAATAAGGCTAGTCCGTTATCAAC |
| **15156** | | GTTGATAACGGACTAGCCTTATTTTAACTTGCTATTTCTAGCTCTAAAACCTAAAGTTCAACAAAGAATTGATCATTTATCTTTCACTGCGGA GAAGTTTCGAACGCCGAAACATGCGCA |
| **15338** | TOM70-6xHIS | CAGCTGAGTCTACTTTGTTAC |
| **15339** | repair amplification | ATCGATGAAGCTATTACATTATTCGAAGAATCC |
| **13807** | URA3 repair | CGGTTTCCTTGAAATTTTTTTGATTCGGTAATCTCCGAACAGAAGGAAGAACGAAGGAAGGGAATCTCGGTCGTAATGATTTCTATAATGAC GAAAAAAAAAAAATTGGAAAGAAAAAGC |
| **13808** | | GCTTTTTCTTTCCAATTTTTTTTTTTTCGTCATTATAGAAATCATTACGACCGAGATTCCCTTCCTTCGTTCTTCCTTCTGTTCGGAGATTACCGA ATCAAAAAAATTTCAAGGAAACCG |
| **15093** | IVT01[2] | GCCGGGAATTTAATACGACTCACTATAGGGAATTTCTACTGTTGTAGATCCGGTTGTGGTATATTTGGAATTTCTACTGTTGTAGAT |
| **15094** | | ATCTACAACAGTAGAAATTCCAAATATACCACAACCGGATCTACAACAGTAGAAATTCCCTATAGTGAGTCGTATTAAATTCCCGGC |
| **18101** | IVT02[2] | GCCGGGAATTTAATACGACTCACTATAGGGAATTTCTACTGTTGTAGATGAGAGCTTCCTCATAACCAAATTTCTACTGTTGTAGAT |
| **18102** | | ATCTACAACAGTAGAAATTTGGTTATGAGGAAGCTCTCATCTACAACAGTAGAAATTCCCTATAGTGAGTCGTATTAAATTCCCGGC |
| **18107** | IVT03[2] | GCCGGGAATTTAATACGACTCACTATAGGGAATTTCTACTGTTGTAGATGAATACCACTTAATTGGTGAATTTCTACTGTTGTAGAT |

| | | |
|---|---|---|
| **18108** | | ATCTACAACAGTAGAAATTCACCAATTAAGTGGTATTCATCTACAACAGTAGAAATTCCCTATAGTGAGTCGTATTAAATTCCCGGC |
| **16745** | IVT04[2] | GCCGGGAATTTAATACGACTCACTATAGGGACTTGAAGATTCTTTAGTGTGTTTTAGAGCTAGAAATAGCAAGT |
| **16476** | | GCACCACCGACTCGGTGCCACTTTTTCAAGTTGATAACGGACTAGCCTTATTTTAACTTGCTATTTCTAGCTCTAAAAC |
| **17690** | IVT-eYFP[2] | GCGAAATTAATACGACTCACTATAGGGAGACC |
| **17691** | | AAAAAACCCCTCAAGACCCGTTTAGAGG |
| **18411** | IVT-Neon[2] | GCCGGGAATTTAATACGACTCACTATAGGGTCTCTAAGGGTGAAGAAGATAACATGG |
| **17385** | | ATGCCGTCTCAGGTCTCAGGATCTTGTACAATTCGTCCATACCC |
| **18413** | IVT-YPET[2] | GCCGGGAATTTAATACGACTCACTATAGGGATGGTCTCCAAGGGCGAAGAATTGTTC |
| **17388** | | ATGCCGTCTCAGGTCTCAGGATTTTATACAATTCATCCATACC |

[1] BsmBI and BsaI recognition sequences are indicated in bold and gRNA sequences are underlined.

[2] Details on IVT construction are indicated under '*In vitro* synthesis of RNA spike-in controls'

#### Table S4

Table S 4. Overview of *in vitro* synthesized RNA controls

| IVT ID | Length [bp] | DNA template | DNA template generation |
|---|---|---|---|
| IVT01 | 60 | 15093 + 15094 | Primer annealing |
| IVT02 | 60 | 18101 + 18102 | Primer annealing |
| IVT03 | 60 | 18107 + 18108 | Primer annealing |
| IVT04 | 102 | 16475 + 16476 | Primer-overlap PCR |
| IVT-eYFP | 1026 | meYFPco-LL-spinach (with 17690 + 17691) | PCR amplification |
| IVT-Neon | 729 | pGGkp304 (with 18411 + 17385) | PCR amplification |
| IVT-YPet | 742 | pGGkp305 (with 18412 + 17388) | PCR amplification |

Table S5

Table S 5. Overview of yields of biomass, mitochondrial biomass and RNA obtained throughout the RNA sequencing process

| | | Ethanol | | | | | Glucose | | | | |
|---|---|---|---|---|---|---|---|---|---|---|---|
| | | Replicate 1 | Replicate 2 | Replicate 3 | Average | St.Dev. | Replicate 1 | Replicate 2 | Replicate 3 | Average | St.dev |
| **Step 1: Mitochondria isolation** | | | | | | | | | | | |
| Culture volume | [mL] | 300.0 | 300.0 | 1200.0 | | | 800.0 | 800.0 | 800.0 | | |
| OD | | 19.0 | 18.6 | 5.8 | | | 5.0 | 5.0 | 5.0 | | |
| Total ODs used | | 5700.0 | 5580.0 | 6996.0 | 6092.0 | 641.1 | 4000.0 | 4000.0 | 4000.0 | 4000.0 | 0.0 |
| Yeast wet weight | [g] | 4.0 | 4.0 | 2.5 | 3.5 | 0.7 | 3.3 | 3.6 | 3.3 | 3.4 | 0.1 |
| **Step 2: RNA isolation** | | | | | | | | | | | |
| RNA yield | [ng/ul] | 115.0 | 119.0 | 129.0 | 121.0 | 5.9 | 862.0 | 726.0 | 632.0 | 740.0 | 94.4 |
| RNA yield in total | [ng] | 4025.0 | 4165.0 | 4515.0 | 4235.0 | 206.1 | 30170.0 | 25410.0 | 22120.0 | 25900.0 | 3304.6 |
| **Step 3: Poly(A)-tailing** | | | | | | | | | | | |
| Input PolyA | [ng] | 3220.0 | 3332.0 | 2805.8 | 3119.3 | 226.3 | 15085.0 | 12705.0 | 11060.0 | 12950.0 | 1652.3 |
| Yield PolyA | [ng/ul] | 90.8 | 71.8 | 34.4 | 65.7 | 23.4 | 232.0 | 159.0 | 222.0 | 204.3 | 32.3 |
| Yield PolyA | [ng] | 1135.0 | 897.5 | 1032.0 | 1021.5 | 97.2 | 6960.0 | 1987.5 | 2775.0 | 3907.5 | 2182.3 |
| **Step 4: Library prep RNA seq** | | | | | | | | | | | |
| Input RNAseq | [ng] | 499.4 | 502.6 | 344.0 | 448.7 | 74.0 | 510.4 | 508.8 | 510.6 | 509.9 | 0.8 |
| Input library | [ng] | 51.0 | 21.0 | 54.6 | 42.2 | 15.1 | 98.0 | 76.0 | 136.0 | 103.3 | 24.8 |
| **Nanopore output** | | | | | | | | | | | |
| RNAseq yield | [gbp] | 0.7 | 0.4 | 0.9 | 0.6 | 0.2 | 1.1 | 1.1 | 1.2 | 1.1 | 0.1 |
| RNAseq yield | [reads] | 1.19E+06 | 4.66E+05 | 1.19E+06 | 9.51E+05 | 3.43E+05 | 1.15E+06 | 1.29E+06 | 1.31E+06 | 1.25E+06 | 6.93E+04 |
| N50 | [bp] | 916.0 | 1347.0 | 1324.0 | 1195.7 | 198.0 | 1394.0 | 1266.0 | 1357.0 | 1339.0 | 53.8 |
| Reads >10kB | [reads] | 10.0 | 8.0 | 8.0 | 8.7 | 0.9 | 8.0 | 6.0 | 11.0 | 8.3 | 2.1 |
| Longest read | [bp] | 50167.0 | 35309.0 | 30844.0 | 38773.3 | 8260.2 | 42954.0 | 16539.0 | 74889.0 | 44794.0 | 23856.8 |

| Nanopore output, filtered for Q >= 7 | | | | | | | | | | | |
|---|---|---|---|---|---|---|---|---|---|---|---|
| RNAseq yield | [gBp] | 0.6 | 0.4 | 0.8 | 0.6 | 0.2 | 1.1 | 1.1 | 1.2 | 1.1 | 0.1 |
| % of total | | 0.9 | 1.0 | 1.0 | | | 1.0 | 1.0 | 1.0 | | |
| RNAseq yield | [reads] | 1.03E+06 | 4.29E+05 | 1.04E+06 | 8.34E+05 | 2.86E+05 | 1.08E+06 | 1.21E+06 | 1.25E+06 | 1.18E+06 | 7.09E+04 |
| % of total | | 0.9 | 0.9 | 0.9 | | | 0.9 | 0.9 | 1.0 | | |
| N50 | [bp] | 939.0 | 1349.0 | 1334.0 | 1207.3 | 189.8 | 1408.0 | 1279.0 | 1367.0 | 1351.3 | 53.8 |
| Read >10kB | [reads] | 0.0 | 0.0 | 0.0 | | | 0.0 | 0.0 | 0.0 | | |
| Longest read | [bp] | 7973.0 | 5494.0 | 7799.0 | 7088.7 | 1129.8 | 8613.0 | 7701.0 | 7869.0 | 8061.0 | 396.3 |

#### Table S6

Table S 6. *De novo* assembly statistics of whole genome sequencing of strains 161-U76 and 161-$I^0$

| | **161-U7** | **161-$I^0$** |
|---|---|---|
| **Total sequence (>500 kb)** | 11.37 Mbp | 11.50 Mbp |
| **Number of contigs (>500 kb)** | 190 | 190 |
| **Largest contig** | 489.45 Kbp | 488.66 Kbp |
| **Average contig length** | 59.82 Kbp | 60.52 Kbp |
| **N50** | 155.57 Kbp | 150.55 Kbp |
| **L50** | 26 | 25 |

#### Table S7

Table S 7. BLAST statistics of the alignment of 161-$I^0$ mtDNA consensus sequence and contigs to the 161-U7 mtDNA consensus sequence

| Description | Max Score | Total Score | Query Cover | E value | Per. ident | Acc. Len | Accession |
|---|---|---|---|---|---|---|---|
| **161-U7 consensus vs 161-I0 consensus** | 7092 | 2.213e+05 | 77% | 0.0 | 100.00% | 67891 | Query_33083 |
| **COX1_161-U7 Vs COX1_161-I0** | 885 | 2853 | 12% | 0.0 | 99.79% | 1605 | Query_43801 |
| **COB_161-U7 Vs COB_161-I0** | 767 | 2120 | 15% | 0.0 | 100.00% | 1158 | Query_31979 |

#### Table S8

Table S 8. Results of running the Dbr1p protein sequence of CEN.PK-type strain (sequence retrieved form SGD) through DeepLoc, numbers indicate the probability of protein targeting to subcellular locations. (16)

| ***Protein_ID*** | ***Localizations*** | ***Signals*** | ***Cytoplasm*** | ***Nucleus*** | ***Extra-cellular*** | ***Cell membrane*** | ***Mitochondrion*** | ***Plastid*** | ***Endoplasmic reticulum*** | ***Lysosome/Vacuole*** | ***Golgi apparatus*** | ***Peroxisome*** |
|---|---|---|---|---|---|---|---|---|---|---|---|---|
| ***Dbr1*** | Cytoplasm \| Nucleus | Nuclear localization signal \| Nuclear export signal | 0.554 | 0.822 | 0.007 | 0.029 | 0.129 | 0.005 | 0.185 | 0.173 | 0.128 | 0.024 |

### Supplementary Methods

#### Cas9 editing of genomic DNA.

Several strains were used in this study but not discussed in the main body of the article (Table S 1). Genomic mutations in the DNA were performed as described in the protocol by (17). URA3 deletion was performed using gRNA plasmid pUDR107 and annealed primers 13807 and 13808 as DNA repair (Table S 2, Table S 3). A 6xHis-tag was added to the mitochondrial Tom70 protein for mitochondrial purification as described by (18), but this mitochondrial isolation method was not used in this study. The 6xHis-tag insertion was performed using plasmid pUDR621 with primers 15155 + 15156 as gRNA and a synthetic *TOM70+6xHis* DNA fragment as repair DNA, ordered as a plasmid at GeneArt (Life technologies, see section 'DNA and RNA sequences') and amplified using primers 15338 + 15339.

#### Western blots

For mitochondrial enrichment analysis by western blot, different fractions of the mitochondrial isolation protocol were kept on ice and normalized to 25-50 mg of biomass in microcentrifuge tubes. The different fractions were spun down and resuspended in 200 µL ice-cold lysis buffer (50 mM HEPES pH 7.5, 150 mM NaCl, 2.5 mM EDTA, 1% v/v Triton X-100, cOmplete mini protease inhibitors (Roche Diagnostics, Rotkreuz, Switzerland)). For whole cells, 100 µL of acid-washed glass beads (425 - 600 micron, Sigma, USA) were added and the suspension was homogenized using a FastPrep-24 (MP Biomedicals, France) in three rounds of 45s shaking at 6.5 m/s and 5 min cooling on ice. For subcellular fractions (homogenate, cytosol, mitochondria, resuspension in lysis buffer and thorough vortexing was sufficient to release proteins. The supernatant was cleared by centrifugation (10.000g for 10 min at 4°C) and protein concentrations were determined using the Quickstart Bradford Protein Assay (Bio-rad Laboratories, Inc, Hercules, CA). 20 µg of total protein was separated on 4-15% Mini-Protean TGX Stain-Free Protein gels (Bio-rad) at 250 V for 20 min. Gels were activated using the stain free protocol on a ChemiDoc MP Imager (Bio-rad) to verify separation of the protein extracts. Proteins were transferred to PVDF membranes using a Turboblotter system (Bio-rad). Successful transfer of proteins was verified using the ChemiDoc MP Imager. Membranes were blocked for 1 hour in TBS + 1% Casein (Blocking Buffer, Bio-rad) and washed once with PBSt (Phosphate saline buffer pH 7.2 with 0.05% Tween20). Washed membranes were incubated overnight, shaken at 4°C, in PBSt with anti-*COX3* antibody, diluted 1 in 1000 (from mouse, #459300, Invitrogen, Waltham, MA). Membranes were washed in PBSt and incubated with the secondary anti-mouse HRP antibody, diluted 1 in 2000 (from goat, Dako Agilent, Santa Clara, CA) for 3 hours at 4°C. Subsequently, membranes were incubated with anti-GAPDH Alexa Fluor 647 antibody for 1 h at room temperature. HRP levels were detected with enhanced chemiluminescence using the Clarity Max Western ECL Substrate (Bio-rad) and chemiluminescence- and fluorescence signals were detected using a ChemiDoc MP Imager (Bio-Rad).

#### Enzyme activity measurements

Protein extracts from whole cells for enzyme activity measurements were prepared as described under 'western blots'. For other fractions of the mitochondrial isolation protocol, fractions were added directly to enzyme activity reaction volumes without any further purification or lysis. Activity of glucose-6-phosphate dehydrogenase (G6PDH) was measured as described by (19), glyceraldehyde-3-phosphate dehydrogenase (GAPDH) was measured according to (20) and activity of cytochrome *c* oxidase was measured as described by (21).

#### Coupling ratio of isolated mitochondria

Coupling ratios of mitochondria was measured by determination of oxygen consumption in 4 mL volume in a stirred chamber at 30 °C and oxygen was measured with a Clark-type oxygen electrode (YSI, Yellow Springs, OH). The chamber was filled with 3,8 mL of Biological Oxygen Measurement (BOM) buffer (0.65 M sorbitol, 25 mM potassium phosphate buffer pH 7.4, 5 mM MgCl). BOM buffer was fully aerated before adding 200 μL of mitochondrial suspension and closing the chamber with the electrode. Then 20μL of succinate (1M pH 7.5) was spiked into the reaction chamber and oxygen consumption rate was measured for a few minutes until at least 10 -20 % of the present oxygen was consumed. Then 10 μL ADP (100mM pH 7.5) was spiked into the chamber and oxygen consumption was measured until oxygen was depleted. The coupling ratio was determined as the ratio between the oxygen consumption rates on succinate prior and after addition of ADP.

#### MNase treatment of mitochondria

For MNase treatment, a protocol by (22) was adapted, together with supplier's instruction of MNase (Thermo Fisher Scientific). Briefly, mitochondrial pellets or mitoplast pellets were resuspended in up to 5 ml MNase buffer (0.25 M sucrose, 10 mM MOPS-KOH, 5 mM $CaCl_2$,) and washed (12.000 × *g*, 10 min, 4°C). Thereafter, mitochondrial or mitoplast pellets were resuspended in 200 - 800 µl MNase buffer, and 1.5 U/µlreaction volume MNase (Thermo Fisher Scientific) was added. Incubation was either performed on ice or at 37 °C for 30 minutes. MNase was inactivated by addition of 0.1 M EGTA solution in nuclease-free water to final concentration of 20 mM and kept for at least 5 minutes on ice. Suspension was centrifuged (12.000 ×g, 10 min, 4°C) and mitochondrial or mitoplast pellets were subject to storage by flash-freezing in liquid nitrogen or RNA extraction.

### DNA and RNA sequences

#### *In vitro* synthesized RNA sequences used in this study

Absence of the *in vitro* sequences from the native, full *S. cerevisiae* nuclear and mitochondrial genome was verified by BLAST. These sequences are of the final *in vitro* transcription product, excluding the long T7 promoter sequence, which was attached by PCR:

```
5'-GCCGGGAATTTAATACGACTCACTATA-3'
```

In the case of IVT-eYFP, also the T7 terminator sequence

```
5'-CTAGCATAACCCCTTGGGGCTCTAAACGGGTCTTGAGGGGTTTTTTG-3'
```

from the DNA template is included, whereas all other IVT sequences do not possess a T7 terminator sequence but are just run-off transcripts.

##### IVT01

```
   1 GGGAAUUUCU ACUGUUGUAG AUCCGGUUGU GGUAUAUUUG GAAUUUCUAC UGUUGUAGAU
```

##### IVT02

```
   1 GGGAAUUUCU ACUGUUGUAG AUGAGAGCUU CCUCAUAACC AAAUUUCUAC UGUUGUAGAU
```

##### IVT03

```
   1 GGGAAUUUCU ACUGUUGUAG AUGAAUACCA CUUAAUUGGU GAAUUUCUAC UGUUGUAGAU
```

##### IVT04

```
   1 GGGACUUGAA GAUUCUUUAG UGUGUUUUAG AGCUAGAAAU AGCAAGUUAA AAUAAGGCUA
  61 GUCCGUUAUC AACUUGAAAA AGUGGCACCG AGUCGGUGGU GC
```

##### IVT-eYFP

```
   1 GGGAAUUUCU ACUGUUGUAG AUCCGGUUGU GGUAUAUUUG GAAUUUCUAC UGUUGUAGAU
  61 UAUGCGGGGU UCUCAUCAUC AUCAUCAUCA UGGUAUGGCU AGCAUGACUG GUGGACAGCA
 121 AAUGGGUCGG GAUCUGUACG ACGAUGACGA UAAGGAUCCG AUGGUUAGCA AAGGCGAAGA
 181 ACUGUUUACG GGCGUGGUGC CGAUUCUGGU GGAACUGGAC GGCGACGUGA ACGGUCACAA
 241 AUUCAGCGUU UCGGGCGAAG GUGAAGGCGA UGCGACCUAU GGUAAACUGA CGCUGAAAUU
 301 UAUUUGCACC ACCGGUAAAC UGCCGGUGCC GUGGCCGACC CUGGUUACCA CGUUUGGUUA
 361 UGGCCUGCAG UGUUUCGCGC GCUACCCGGA UCAUAUGAAA CAACACGACU UUUUCAAAUC
 421 UGCCAUGCCG GAAGGUUAUG UGCAGGAACG UACGAUUUUC UUUAAAGAUG ACGGCAACUA
 481 CAAAACCCGC GCAGAAGUCA AAUUUGAAGG UGAUACGCUG GUGAACCGUA UUGAACUGAA
 541 AGGCAUCGAU UUCAAAGAAG ACGGUAAUAU CCUGGGCCAU AAACUGGAAU ACAACUACAA
 601 CUCCCACAAC GUUUACAUCA UGGCAGAUAA ACAGAAAAAC GGUAUCAAAG UCAACUUCAA
 661 AAUCCGCCAU AACAUCGAAG AUGGCUCAGU GCAACUGGCU GACCACUACC AGCAAAACAC
 721 CCCGAUCGGU GAUGGCCCGG UUCUGCUGCC GGACAAUCAU UAUCUGAGCU ACCAGUCUAA
 781 ACUGAGUAAA GAUCCGAACG AAAAACGUGA CCACAUGGUC CUGCUGGAAU UUGUGACGGC
 841 GGCUGGUAUU ACGCUGGGCA UGGAUGAACU GUAUAAAUGA AAGCUUCCCG GGAAAGUAUA
 901 UAUGAGUAAA GAUAUCGACG CAACUGAAUG AAAUGGUGAA GGACGGGUCC AGGUGUGGCU
 961 GCUUCGGCAG UGCAGCUUGU UGAGUAGAGU GUGAGCUCCG UAACUAGUCG CGUCGAUAUC
1021 CCCGGG
```

#### IVT-Neon

```
  1 GGGUCUCUAA GGGUGAAGAA GAUAACAUGG CUUCUUUGCC AGCCACCCAC GAGUUGCACA
 61 UCUUCGGUUC CAUUAACGGU GUCGACUUCG AUAUGGUCGG UCAAGGUACC GGUAACCCAA
121 ACGACGGUUA CGAAGAAUUG AACUUGAAGU CUACCAAGGG CGACUUGCAG UUCUCCCCUU
181 GGAUUUUGGU UCCACACAUC GGUUACGGUU UCCACCAAUA CUUGCCAUAC CCAGACGGUA
241 UGUCCCCAUU CCAAGCCGCC AUGGUUGACG GUUCUGGUUA CCAAGUCCAC AGAACCAUGC
301 AAUUCGAAGA CGGUGCUUCC UUGACCGUGA ACUACAGAUA CACCUACGAA GGUUCUCACA
361 UCAAGGGCGA AGCCCAAGUU AAGGGUACCG GUUUCCCUGC UGACGGUCCA GUCAUGACUA
421 ACUCCUUGAC CGCUGCUGAC UGGUGUCGUU CCAAGAAGAC CUACCCAAAC GAUAAGACCA
481 UCAUUUCCAC UUUCAAGUGG UCCUACACCA CCGGUAACGG CAAGAGAUAC CGCUCUACCG
541 CUCGUACCAC CUACACCUUC GCCAAGCCAA UGGCUGCCAA CUACUUGAAG AACCAGCCUA
601 UGUACGUCUU CCGUAAGACC GAGCUGAAGC AUUCCAAGAC UGAAUUGAAC UUCAAGGAAU
661 GGCAAAAGGC UUUCACCGAC GUCAUGGGUA UGGACGAAUU GUACAAGAUC CUGAGACCUG
721 AGACGGCAU
```

#### IVT-YPet

```
  1 GGGAUGGUCU CCAAGGGCGA AGAAUUGUUC ACCGGUGUGG UCCCAAUCUU GGUCGAGUUG
 61 GACGGUGACG UCAACGGUCA CAAGUUCUCU GUUUCCGGUG AAGGUGAAGG AGACGCCACC
121 UACGGUAAAU UGACCUUGAA GUUGUUGUGC ACCACCGGCA AGCUGCCAGU UCCAUGGCCA
181 ACAUUGGUCA CCACUUUAGG UUAUGGUUUA CAAUGUUUUG CUAGAUAUCC AGAUCAUAUG
241 AAACAACAUG AUUUCUUCAA AUCUGCUAUG CCAGAAGGUU AUGUUCAAGA AAGAACUAUA
301 UUUUUCAAAG AUGAUGGUAA UUAUAAAACU AGAGCUGAAG UUAAAUUUGA AGGAGAUACU
361 UUAGUUAAUA GAAUUGAAUU GAAAGGUAUU GAUUUUAAAG AAGAUGGUAA UAUUUUAGGU
421 CAUAAAUUAG AAUAUAAUUA UAAUUCUCAU AAUGUUUAUA UUACAGCUGA UAAACAAAAG
481 AAUGGUAUUA AGGCUAAUUU UAAAAUUAGA CAUAAUAUUG AAGAUGGUGG UGUUCAAUUA
541 GCUGAUCAUU AUCAACAAAA UACUCCUAUU GGUGACGGUC CAGUUUUAUU ACCUGAUAAU
601 CAUUAUUUGU CAUAUCAAUC UAAAUUGUCA AAAGAUCCAA AUGAAAAAAG AGAUCAUAUG
661 GUUUUGUUAG AAUUUUUGAC UGCAGCUGGU AUUACUUUAG GUAUGGAUGA AUUGUAUAAA
721 AUCCUGAGAC CUGAGACGGC AU
```

### Synthetic DNA sequences used in this study

#### PreSU9 mitochondrial targeting signal

```
  1 ATGGCTTCTA CTAGAGTTTT GGCTTCTAGA TTGGCTTCTC AAATGGCTGC TTCTGCTAAG
 61 GTTGCTAGAC CAGCTGTTAG AGTTGCTCAA GTTTCTAAGA GAACTATCCA AACTGGTTCT
121 CCATTGCAAA CTTTGAAGAG AACTCAAATG ACTTCTATCG TTAACGCTAC TACTAGACAA
181 GCTTTCCAAA AGAGAGCT
```

1. Hinkle, P.C. (2005) P/O ratios of mitochondrial oxidative phosphorylation. *Biochimica et Biophysica Acta (BBA) - Bioenergetics*, **1706**, 1-11.
2. Turk, E.M., Das, V., Seibert, R.D. and Andrulis, E.D.J.P.O. (2013) The mitochondrial RNA landscape of Saccharomyces cerevisiae. **8**.
3. Di Bartolomeo, F., Malina, C., Campbell, K., Mormino, M., Fuchs, J., Vorontsov, E., Gustafsson, C.M. and Nielsen, J. (2020) Absolute yeast mitochondrial proteome quantification reveals trade-off between biosynthesis and energy generation during diauxic shift. *Proceedings of the National Academy of Sciences*.
4. Mitchell, S.F., Jain, S., She, M. and Parker, R. (2013) Global analysis of yeast mRNPs. *Nature Structural & Molecular Biology*, **20**, 127-133.
5. Tsvetanova, N.G., Klass, D.M., Salzman, J. and Brown, P.O. (2010) Proteome-Wide Search Reveals Unexpected RNA-Binding Proteins in *Saccharomyces cerevisiae*. *PLoS One*, **5**, e12671.
6. Di Bartolomeo, F., Malina, C., Campbell, K., Mormino, M., Fuchs, J., Vorontsov, E., Gustafsson, C.M. and Nielsen, J. (2020) Absolute yeast mitochondrial proteome quantification reveals trade-off between biosynthesis and energy generation during diauxic shift. *Proceedings of the National Academy of Sciences*, **117**, 7524-7535.
7. Yang, J., Mohr, G., Perlman, P.S. and Lambowitz, A.M. (1998) Group II intron mobility in yeast mitochondria: target DNA-primed reverse transcription activity of ai1 and reverse splicing into DNA transposition sites *in vitro*. *Journal of Molecular Biology*, **282**, 505-523.
8. Schmelzer, C. and Schweyen, R.J. (1986) Self-splicing of group II introns *in vitro*: Mapping of the branch point and mutational inhibition of lariat formation. *Cell*, **46**, 557-565.
9. Randazzo, P., Bennis, N.X., Daran, J.-M. and Daran-Lapujade, P. (2021) gEL DNA: A cloning- and polymerase chain reaction–free method for CRISPR-based multiplexed genome editing. *The CRISPR Journal*, **4**, 896-913.
10. Bouwknegt, J., Koster, C.C., Vos, A.M., Ortiz-Merino, R.A., Wassink, M., Luttik, M.A.H., van den Broek, M., Hagedoorn, P.L. and Pronk, J.T. (2021) Class-II dihydroorotate dehydrogenases from three phylogenetically distant fungi support anaerobic pyrimidine biosynthesis. *Fungal Biology and Biotechnology*, **8**, 10.
11. Boonekamp, F.J., Knibbe, E., Vieira-Lara, M.A., Wijsman, M., Luttik, M.A.H., van Eunen, K., Ridder, M.d., Bron, R., Almonacid Suarez, A.M., van Rijn, P. *et al.* (2022) Full humanization of the glycolytic pathway in *Saccharomyces cerevisiae*. *Cell Reports*, **39**, 111010.
12. Lee, M.E., DeLoache, W.C., Cervantes, B. and Dueber, J.E. (2015) A highly characterized yeast toolkit for modular, multipart assembly. *ACS synthetic biology*, **4**, 975-986.
13. Botman, D., de Groot, D.H., Schmidt, P., Goedhart, J. and Teusink, B. (2019) *In vivo* characterisation of fluorescent proteins in budding yeast. *Scientific reports*, **9**, 2234.
14. Gorter de Vries, A.R., de Groot, P.A., van den Broek, M. and Daran, J.-M.G. (2017) CRISPR-Cas9 mediated gene deletions in lager yeast *Saccharomyces pastorianus*. *Microbial Cell Factories*, **16**, 222.
15. Doerr, A., De Reus, E., Van Nies, P., Van der Haar, M., Wei, K., Kattan, J., Wahl, A. and Danelon, C. (2019) Modelling cell-free RNA and protein synthesis with minimal systems. *Physical biology*, **16**, 025001.
